# From FODMAPs to prebiotic candidates: enzymatic transglycosylation of raffinose oligosaccharides towards new mixed-linkage oligosaccharides

**DOI:** 10.64898/2026.06.09.731070

**Authors:** Philipp Garbers, Gordon Jacob Boehlich, Birgitte Zeuner, Jane Wittrup Agger, Bjørge Westereng

## Abstract

Raffinose family oligosaccharides (RFOs) are abundant in side streams from food and feed production from legumes, and the transition to plant-based diets increases the volume of such side streams. RFOs in the diet tend to have negative impacts on the consumer’s gut (e.g., nausea, bloating, diarrhoea), and in many ways, RFOs are comparable to lactose as a side stream from the dairy industry and symptoms associated with lactose intolerance. On the contrary, galactooligosaccharides (GOS) are recognized as prebiotics, and in this study we used a β-galactosidase from *Niallia circulans* to produce potential prebiotics from RFOs (acceptors) and lactose (donor), which we hypothesized to have a lower fermentability than unmodified RFOs. The transglycosylation reactions resulted in RFO-based α-β-GOS, with NMR characterization showing (β1-4) galactosylations on the non-reducing galactose end of RFOs as the major product. In reactions with RFOs, the characteristics were comparable to reactions with lactose alone and the new α-β-GOS products made up the largest fraction (by weight). A screening of 11 relevant gut and food microbe strains revealed that the gut commensal *Bacteroides ovatus* metabolised these modified oligosaccharides for growth whereas other strains grew only after adaption and others did not use them at all. This implies that mixed-linkage α-β-GOS are less fermentable by some microbes compared to raffinose, while other (beneficial) bacteria can still ferment them. The enzymatic synthesis established here is an interesting approach to upgrade abundant food side streams towards new prebiotics in a world where functional foods and food waste reduction receive increasing attention.

**Graphical abstract:** 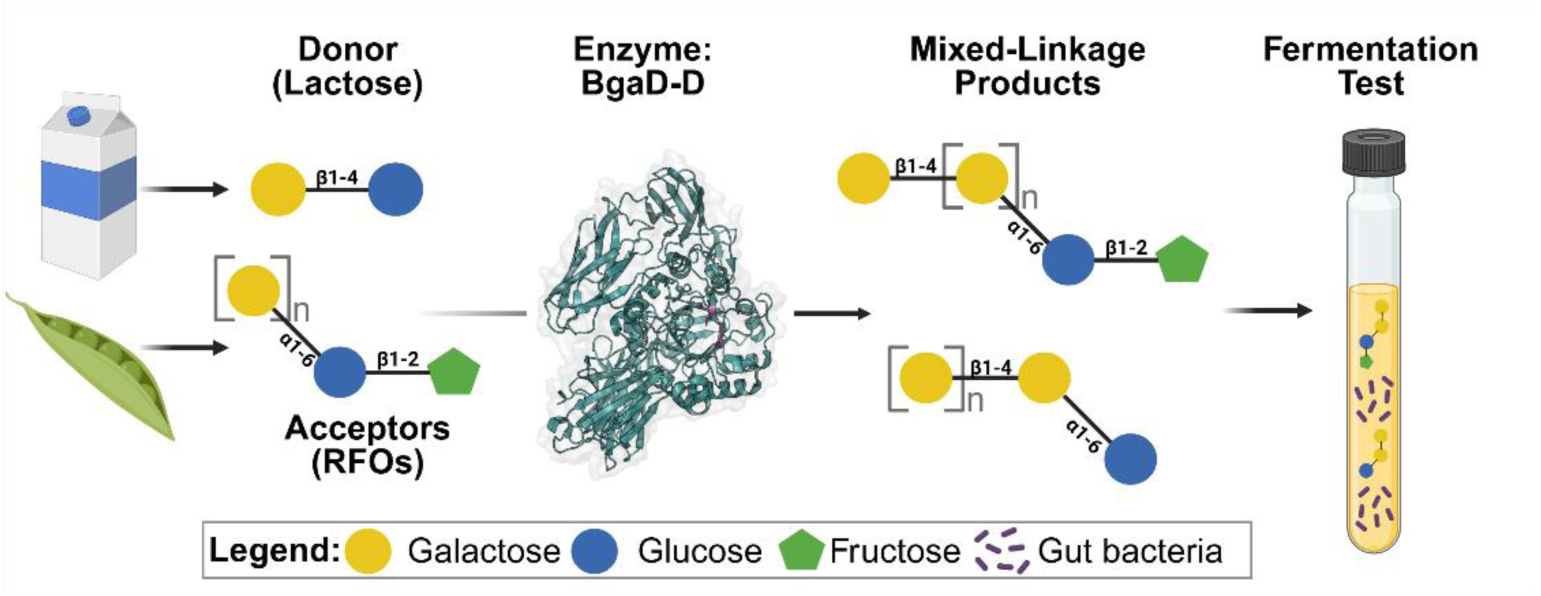

## 1. Introduction

Raffinose family oligosaccharides (RFOs, Figure 1) are abundant in starch- and oil-rich legumes, such as peas (Daveby et al., 2006), faba beans (Njoumi et al., 2019) and soybeans (Kim et al., 2003). Since the human genome does not encode genes for enzymes degrading RFOs, like α-galactosidases and invertases, humans are unable to digest these oligosaccharides. As a result, RFOs pass through to the lower intestine where bacteria from the gut microbiome digest them and RFOs are thus considered fermentable oligo-, di-, and monosaccharides and polyols (FODMAPs) (Gibson & Shepherd, 2005). This can lead to the production of gas, causing flatulence and discomfort, and in more severer cases lead to nausea and diarrhoea (Elango et al., 2022). But RFOs are also easily fermented by a range of food-related lactic acid bacteria (Garbers et al., 2025) and beneficial bacteria such as *Bifidobacterium* (van Zanten et al., 2012). While the effects of RFOs are not consistent in all individuals and the definition of FODMAPs remain open for debate, high levels of RFOs in diets are problematic (Elango 2022, Gibson 2020). Yet, RFOs received GRAS status in the USA (Kim et al., 2003) and are acknowledged as a prebiotic in Japan (Elango et al., 2022) which eases implementation in future functional foods and prebiotics. In the food industry, RFOs occur in large amounts as part of the leftovers from soybean processing (Kim et al., 2003) (industrial plants process up to 22.000 t per day, with a reported content of approximately 5% RFOs (Demarco & Gibon, 2020; Zhang et al., 2019). RFOs are also important constituents in other legume products, most notably chickpea, aquafaba and pea and faba bean protein concentrates (Huang et al., 2024; Saldanha do Carmo et al., 2022), that are often used in the production of so-called meat alternatives and other processed plant-based foods (Saldanha do Carmo et al., 2021). However, RFOs can be extracted from the protein-rich ingredients through water-based processes (Garbers et al., 2026) or by utilizing ethanol (Gulewicz et al., 2000). Presumably, such extraction processes are already in place in the production of protein isolates from faba bean for example, as these isolates have reduced RFO contents (Bhatty & Christison, 1984; Vogelsang-O’Dwyer et al., 2020). RFOs therefore make up a notable amount of food side streams and are routinely ingested through consumption of pulses.

**Figure 1:**
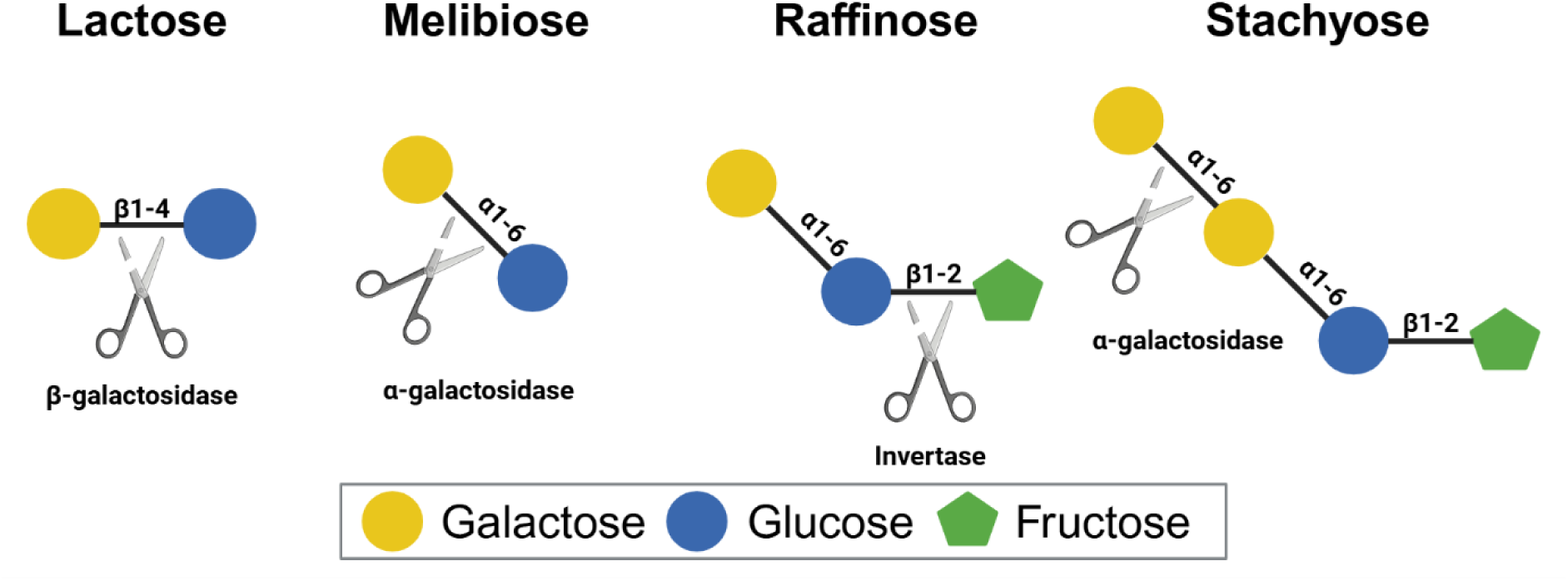
The structure of raffinose family oligosaccharides and lactose represented in accordance with Essentials of Glycobiology, 4^th^ Edition (Varki et al., 2022) in addition to the required enzymatic actions for hydrolysis of these oligosaccharides.

Another common side stream in the food industry is lactose (Figure 1), which is available from dairy processing, for example in whey (Jameson et al., 2021) and can lead to FODMAP-like effects in consumers with intolerance (Swagerty et al., 2002). Lactose is however commonly used as building block for enzymatic and whole-cell synthesis of galactooligosaccharides (GOS) (Gänzle, 2012) and human milk oligosaccharides (HMOs) (Zeuner et al., 2016). Both are considered prebiotic, and HMOs are even regarded as essential for the development of a healthy infant gut microbiome and have been shown to especially promote growth of *Bifidobacterium sp.* (Barile & Rastall, 2013). GOS are commonly produced by microbial enzymes rather than chemical synthesis (Wang et al., 2024) through β-galactosidases with a strong preference for transglycosylation (Barile & Rastall, 2013). . One frequently used enzyme, in both industry and research, is the β-galactosidase BgaD from *Niallia circulans* ATCC 31382 (formerly *Bacillus circulans*). The enzyme belongs to glycoside hydrolase family 2, subfamily 1 (Lebreton et al., 2025), uses a wide range of acceptors (Farkas et al., 2003) and can operate at elevated temperatures (Zeuner et al., 2016). Commercially established GOS products obtained with BgaD contain a variety of oligosaccharide structures (> 40) with different lengths up to 8 monomeric units (van Leeuwen et al., 2014). This broad product profile stems from the mixture of isoforms of BgaD in most enzyme preparations, and of particular interest is the BgaD-D isoform as the majority of products are of (β1-4) type (Nekvasilova et al., 2022; Yin et al., 2017a; Yin et al., 2017b), creating a smaller set of products compared to the mixtures of all isoforms (van Leeuwen et al., 2014). Additionally, it has the least structural obstructions for creating larger oligosaccharides (Hovorkova et al., 2024). Raffinose is among the suitable acceptors reported for β-galactosidases from *N. circulans* (Farkas et al., 2003), however, the potential of stachyose, verbascose or melibiose has not been tested and only single transgalactosylations to raffinose were demonstrated.

In this study we investigated the synthesis of new oligosaccharides from RFOs and lactose, with a combination of α- and β-linked galactose, glucose and fructose units (α-β-GOS). The transglycosylation behaviour in the presence of different RFO acceptors, acceptor-donor and donor-enzyme ratios were investigated and the products were structurally characterized through Nuclear Magnetic Resonance (NMR) spectroscopy to allow detailed structural elucidation of the products. It is a underlying hypothesis that mixed-linkage oligosaccharides can mitigate FODMAP effects, and that the increased complexity might also result in an increased selectivity by fermenting organisms and eventually result in a prebiotic effect. To evaluate the effect of the modifications, we pre-screened a variety of gut and food bacteria for growth with unpurified transglycosylation products. Additionally, the fermentability of two purified and characterized oligosaccharides was assessed in detail for three bacterial strains of interest.

## 2. Material & Methods

### 2.1. Materials

All materials and analytical standards were purchased from either Sigma-Aldrich (Merck, Darmstadt, Germany) or VWR (Avantor, Radnor, Pennsylvania, United States) except for stachyose, which was purchased from BioSynth (Staad, St. Gallen, Switzerland). Growth media and its components were purchased at purities suitable for microbiology, whereas standards for analytics were of the best available purity like “analytical standard” (monosaccharides, lactose) or at least ≥ 98 % purity (e.g., melibiose, raffinose, stachyose).

The RFO extract was produced as previously described (Garbers et al., 2026) from pea protein concentrate in a pilot scale biorefining process through water-based extraction and purification. In order to reduce the content of sucrose, a batch of RFO extract was precipitated in 90 % v/v ethanol as described in literature (Gulewicz et al., 2000), while sucrose mainly remains in solution. The precipitated RFOs were then separated by centrifugation and dried before being resuspended in water for reactions.

The β-galactosidase BgaD-D from *N. circulans* was produced as described previously (Zeuner et al., 2016) in *E.coli* BL21 star. For larger scale productions (4 L volume), kanamycin resistance was replaced by ampicillin resistance and cells were only pre-cultured in lysogeny broth (LB) before grown and induced in terrific broth (TB) in multiple 1 L Infors HT Minifors 2 fermenters (Infors AG, Bottmingen, Switzerland) with aeration (1 L air/min) and titration (H_2_SO_4_, NaOH). Proteins were purified by IMAC chromatography (His-Tag) on a BioRad chromatography system (Hercules, CA, United States) and confirmed by SDS-PAGE. Protein concentrations were determined by absorbance a 280 nm with an extinction coefficient for BgaD-D of 175,000 M^−1^ cm^−1^. Aliquots of enzyme were frozen at -20°C with 10 % v/v glycerol in the buffer.

Additionally, invertase (GH32) from *Saccharomyces cerevisiae* was purchased from Sigma-Aldrich and an α-galactosidase (GH36) from *Aspergillus niger* was purchased from Megazyme (Wicklow, Ireland). These were used to remove the terminal α-galactose and fructose from raffinose and its derivatives.

### 2.2. Reactions

All reactions were performed in 1.5 mL and 2.0 mL reaction tubes and reaction conditions were controlled in a Thermomixer with a ThermoCap (Eppendorf, Hamburg, Germany). Temperatures were kept at 50°C for BgaD-D and shaking was always set to 1000 rpm. Reactions were terminated through heating the samples to 100°C for 5-10 min. Reactions were performed at different enzyme and substrate concentrations as specified in the according results sections and monitored for up to 180 min. Invertase & α-galactosidase reactions were performed at 37°C for at least 60 min.

### 2.3. Reduction

Of reducing ends was performed as previously described (Westereng et al., 2020): 10 µL of sample were alkalized with 10 µL 25 mM NaOH and then 65 µL aqueous NaBD_4_ solution (10.77 g/L) was added. The reduction was performed for ≥6 hours before stopping the reaction by slightly acidifying the mixture (15 µL of 25 mM CH_3_COOH). Chromatography (HPAEC-PAD, section 2.5.1) and mass spectrometry (MALDI-TOF, section 2.5.2) was then performed as is.

### 2.4. Microbial growth experiments

A complete list of all strains and some of their enzymatic tools according to literature is given below (Table 1) and recipes of all prepared growth media are available in the Supplementary Materials. Strains of different lactic acid bacteria (LAB) were chosen from a previous study based on their growth in presence of RFOs and cultures were aerobically prepared as described therein with either deMan-Rogosa-Sharpe (MRS) or M17 media (Garbers et al., 2025). Strains of *Bacteroides* (La Rosa et al., 2019; Leivers et al., 2022; Lindstad et al., 2021), *Roseburia* (La Rosa et al., 2019) and *Faecalibacterium* (Lindstad et al., 2021) were chosen due to their importance in the gut and their vast toolbox of carbohydrate active enzymes (CAZymes) (Lombard et al., 2014). These strains were cultured in an anaerobic cabinet (Whitley A85 TG Workstation; Don Whitley, UK) with 85 % N_2_, 5 % CO_2_ and 10 % H_2_ gas and grown at 37°C, as previously described (Leivers et al., 2022). All pre-cultures were grown with 5 g/L glucose before being transferred to media containing 5 g/L of different carbon sources. Carbon sources were either pure compounds or reaction mixtures after boiling (see section 2.2). Both aerobic and anaerobic cultures were inoculated to the same degree by diluting precultures and adjusting inoculation volume according to the total volume to facilitate similar growth curves under all conditions. Growth was assessed by means of turbidity through absorbance at 600 nm (OD_600_) up to 72 hours and all experiments were performed in triplicates. All anaerobic fermentations were carried out in culture tubes. Turbidity was manually measured with a portable spectrophotometer at different time points. The instrument was blanked against a control tube with media without carbon source. Aerobic fermentations on the other hand were conducted in 96-well plates, without shaking, each well filled with 150 µL of media and the culture density was measured with a plate-reader automatically every 15 mins over 48 hours. The pathway of the plate filled with 150 µL was estimated to be 0.42 cm compared to 1 cm with culture tubes. Measurements have been corrected accordingly with this factor, and a cu-inoculated blank has been subtracted from the samples. Samples for chromatography were taken at the end of the experiment and at some intermediate timepoints, sterile filtered and compared to the media without any addition of microorganisms. If analysis was performed at a later stage, samples were frozen at -20°C until preparation. Additionally, pH was roughly measured at the end point with pH-strips and compared to pure media.

**Table 1:** List of all tested bacterial strains as well as number relevant glycoside hydrolases (GH) in the strains genomes according to *cazy.org database (Lombard et al., 2014), **DSMZ/BacDrive datasheet (positive = substrate proven as carbon source and/or as fermentable) (BacDrive, 2025; Schober et al., 2025), ***internal data. Empty fields indicate no information. All strains are commercially available, except for *L. pentosus* KW1 / KW2 (Wiull et al., 2024) and *L. cremoris* 121 (Garbers et al., 2025), which are in-house strains.

| Strain | β-galactosidase | | α-galactosidase | | | | Invert-ase |
| --- | --- | --- | --- | --- | --- | --- | --- |
|  | GH2 | GH42 | GH27 | GH31 | GH36 | GH97 | GH32 |
| *Faecalibacterium prausnitzii* SL3/3* | 7 | 0 | 0 | 1 | 2 | 0 | 0 |
| *Roseburia intestinalis* L1-82* | 10/11 | 2 | 2 | 4 | 3 | 0 | 1 |
| *Bacteroides cellulosilyticus* DSMZ 14838** | Pos. | Pos. | Pos. |  | Pos. |  | Pos. |
| *Bacteroides ovatus* ATCC 8483* | 37 | 1 | 3 | 12 | 3 | 12 | 2 |
| *Bacteroides thetaiotaomicron* 7330* | 31 | 1 | 4 | 6 | 3 | 11 | 4 |
| *Lactocaseibacillus rhamnosus* GG* | 2/3 | 0 | 0 | 1 | 2 | 0 | 1 |
| *Levilactobacillus brevis* BSO 464* | 3 | 1 | 0 | 2 | 1 | 0 | 0 |
| *Lentilactobacillus buchneri* subspecies<br>*silagei* CD034* | 3 | 0 | 1 | 1 | 1 | 0 | 0 |
| *Lactiplantibacillus pentosus* KW1* | 2 | 0 | 0 | 1 | 1 | 0 | 1 |
| *Lactiplantibacillus pentosus* KW2* | 2 | 0 | 0 | 1 | 1 | 0 | 1 |
| *Lactococcus lactis* subspecies *cremoris*<br>121*** | 0 | 0 | 1 | 1 | 0 | 0 | 0 |

### 2.5. Chemical analysis

#### 2.5.1. High Performance Anion Exchange Chromatography with Pulsed Amperometric Detection (HPAEC-PAD)

Quantitative analysis of saccharides was performed on a Dionex ICS6000 system (Thermo Fisher Scientific, Waltham, Massachusetts, USA) equipped with a CarboPac PA210 Fast 4µm 2x250mm column (Thermo Fisher Scientific) with the same methods as previously described (Garbers et al., 2025).

Relevant analytes were identified and quantified by comparison with external standards and calibration curves (to the extent possible). Data analysis was performed with the Chromeleon 7.2.9 software (Thermo Fisher Scientific).

Based on HPAEC results, the following definitions and calculations were used in Figure 2:

1. Conversion is the % of lactose that has been degraded compared to the starting concentration at 0 min;
2. Hydrolysis of lactose is defined as free galactose concentrations;
3. Transglycosylation towards RFOs are defined as reduction in RFO concentration;
4. Transglycosylation towards lactose is defined as the difference between reduction in lactose and free glucose concentrations.
5. Additional transglycosylations to any acceptor (e.g., multiple to the same molecule) are defined as the reduction of lactose minus (1), (2) and (3).

**Figure 2:**
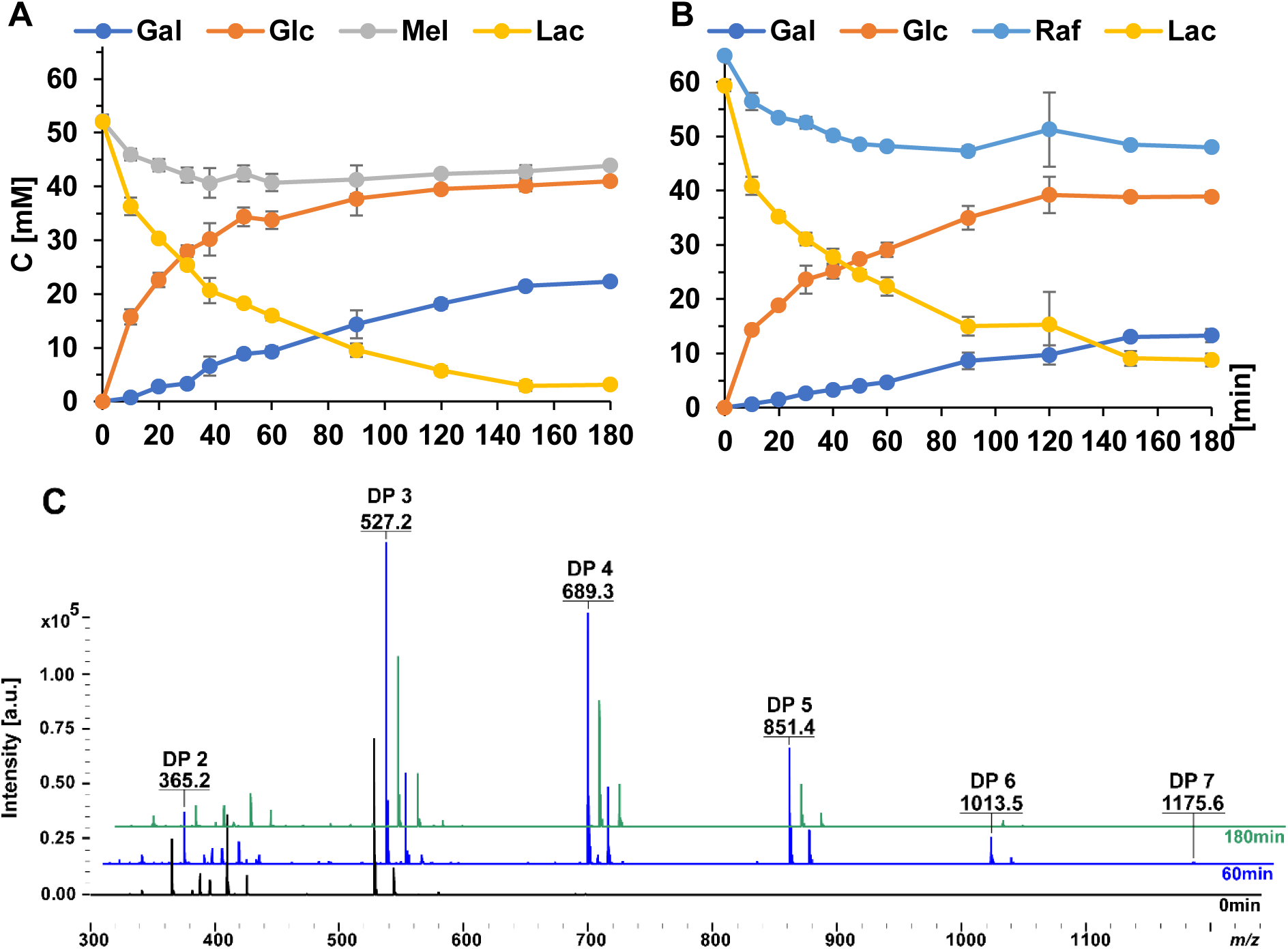
Time course reactions (0-180 minutes) with 0.25 µM β-galactosidase (BgaD-D), equimolar mixture of lactose and either **A:** melibiose or **B:** raffinose (50mM each) at 50°C, symbols: Gal, galactose; Glc, glucose; Mel, melibiose; Lac, lactose; Raf, raffinose, [min], minutes. Concentrations of acceptors, donors and products were measured by HPAEC-PAD. **C:** MALDI-TOF spectra of reactions at 50°C with BgaD-D (0.25 µM) raffinose (50mM), lactose (50mM) after 0 (black), 60 (blue) and 180 min (green). Spectra are offset by 10 *m/z*.

#### 2.5.2. Hydrophilic Interaction Liquid Chromatography (HILIC)

At analytical scale a Waters XBridge Amide 3.5µm 4.6x250mm column and a Thermo Fisher Scientific UltiMate 3000 chromatography system with a charged aerosol detector (CAD; esa Corona ultra, Thermo Fisher Scientific) was used. Eluents were purified water with 0.1 % v/v ammonium and 65 % acetonitrile at a flow of 1 mL/min at 35 °C. This setup was used to analyse fermentations of purified oligosaccharides with injection volumes of 10 µL.

Preparative HILIC was performed to purify oligosaccharides after transglycosylation reactions at a suitable product scale to test them for fermentations as well as for structural characterization with nuclear magnetic resonance (see 2.5.4). The analytical column was replaced by a preparative Waters XBridge BEH Prep OBD Amide 5µm 30x250mm Column (Waters Corporation, Framingham, Massachusetts, United States) with a flow of 40 mL/min at room temperature was but otherwise same eluents on an Agilent Technologies 1260 Infinity system (Santa Clara, CA, United States) equipped with refractive index (RI) ultra violet light (UV, 280 nm) detectors. Injections were 900 µL with 225 mg material.

#### 2.5.3. Matrix Assisted Laser Desorption Ionization Time of Flight (MALDI TOF) mass spectrometry

Samples of 1 µL were spotted with 2 µL of matrix (9 mg/mL dihydroxybenzoic acid in 30% acetonitrile) and analyzed on a Bruker ultraFlextreme (Bruker Daltonics, Bremen, Germany). Data acquisition and analysis were performed in flexControl 3.4 and flexAnalysis 3.4 (Bruker Daltonics).

#### 2.5.4. Nuclear Magnetic Resonance (NMR) Spectrometry

All materials analyzed by NMR were dissolved in D_2_O and lyophilized to minimize the HDO signal. Finally, samples were dissolved in 0.5 mL D_2_O containing 0.05 wt. % 3 (trimethylsilyl)propionic-2,2,3,3-*d*_4_ acid, sodium salt. The samples were measured using 5 mm Norell® Sample Vault Series™ NMR tubes. ^1^H-NMR and ^13^C-NMR spectra were recorded at 298 K using a Bruker AVANCE III HD 400 MHz instrument (Bruker BioSpin, Ettlingen, Germany) equipped with BBFO room temperature probe. Chemical shifts are reported in ppm with Na-Trimethylsilylpropanoate-*d*_4_ (δ ^1^H/^13^C = 0.00 ppm) as internal standard. Multiplicities are reported as follows: s = singlet, d = doublet, t = triplet q = quartet, m = multiplet. Coupling constants (*J*) are reported in Hz. The following pulse programs where used: “zg30” for ^1^H, “deptqgpsp.2” for DEPTQ-^13^C, “cosygpmfppqf” for ^1^H-^1^H COSY, “dipsi2gpphzs” for TOCSY “hsqcetgpsisp2.2” for HSQC, “h2bcetgpl3” for H2BC, “hsqcdietgpsisp.2” for HSQC-TOCSY and “hmbcetgpl3nd” for HMBC.

### 2.6. Statistical analysis

Significant differences between samples (Figure 2) were first assessed for homogeneity of variances via Levene’s test, and data tested for following a normal distribution via inspection of Q-Q residual plots and the Shapiro-Wilk test. As the data were not normally distributed, the Kruskal-Wallis test was performed to test for significant differences among the same levels of substrates used in the experiment. All statistical analyses were performed in R 4.3.2.

## 3. Results & Discussion

### 3.1. BgaD-D reactions

#### 3.1.1 Time courses

To determine the ideal reaction time and demonstrate activity of BgaD-D on an equimolar mixture of either melibiose or raffinose with lactose, the reaction was monitored for 3 hours with an enzyme concentration of 0.25 µM and 50 mM of each substrate (Figure 2). In the presence of melibiose, about half (51.3 %) of the lactose was converted within the first 30 min (Figure 2 A), before the rate of the reaction slowed down and the conversion reached a maximum of 94.5 % at 150 min. Glucose release followed the inverted trend of lactose depletion. Galactose on the other hand was released slowly in a linear fashion compared to glucose. The difference in galactose and glucose release is equal to the amount of transglycosylation occurring. While some of the transglycosylation took place with lactose as acceptor, a part of the transglycosylation seemingly used melibiose as acceptor, as the melibiose concentration also decreased during the reaction with a maximum decrease of 22.1 % after 30 min. Notably, control reactions confirmed that BgaD-D is not active on pure melibiose (Supplementary Figure 1), implying that the reduction in melibiose is a result of transglycosylation rather than degradation. New peaks appeared in chromatograms when melibiose was present, compared to BgaD-D reactions with lactose alone (Suppl. Figure 2), corroborating that melibiose is functioning as an acceptor. Lactose conversion appears higher than glucose release, with a difference of up to 9 mM, and this likely stems from transglycosylation with lactose as the acceptor. The almost linear galactose increase observed until 150 min could stem from hydrolysis of lactose as well as of the transglycosylation products. The slight increase in melibiose after ≥60 min of reaction (Figure 2 A) suggests product hydrolysis, a type of event commonly referred to as secondary hydrolysis which causes a temporary product maximum (Zeuner et al., 2019). This was also reported for reactions with only lactose and BgaD-D (Warmerdam et al., 2013). Hydrolysis of transglycosylation products is also visible in MALDI-TOF MS, where the maximum detected length of oligosaccharides (DP 7) decreases to DP 6 after 90 min and even further to DP 5 after 180 min (Table 2). The longest detected oligosaccharides (DP 7) were visible after 10 min of reaction with melibiose as acceptor. In a commercially available GOS mixture produced with all four isoforms of BgaD-D, DP up to 8 was previously reported, yet oligosaccharides of DP6 or higher only made up 0.5 % of the total products (van Leeuwen et al., 2014). While glucose release decreased over time, galactose release remained relatively constant until 150 min indicating a relative increase of hydrolysis compared to transglycosylation towards the end; at lower lactose concentrations hydrolysis has been shown to be higher (Warmerdam et al., 2013). Additionally, affinity between BgaD-D and its transglycosylation products seems to be lower than for lactose, as demonstrated by exposure of the boiled reaction mixture to a fresh batch of enzyme, which did not lead to full degradation of elongated products (Suppl. Figure 3). Similar observations have been made for some α-galactosidases where hydrolysis activity decreases with increasing RFO lengths (Shin et al., 2020). Warmerdam et al. (2013) showed, however, that BgaD-D can degrade 91 % its own product (4-galactosyl-lactose, DP 3) within an hour, but not completely to monosaccharides. This aligns with the results obtained in our study (Suppl. Figure 3).

**Table 2:** Maximum length of oligosaccharides detected by MALDI-TOF at different time points [min] over the course of three hours for reactions between equimolar concentrations of lactose and melibiose or raffinose respectively. Conditions: 50°C, 1000 rpm, 0.25 µM BgaD-D, 50 mM initial lactose.

| Time | 0 | 10 | 20 | 30 | 40 | 50 | 60 | 90 | 120 | 150 | 180 | [min] |
| --- | --- | --- | --- | --- | --- | --- | --- | --- | --- | --- | --- | --- |
| **Melibiose** | 2 | 7 | 7 | 7 | 7 | 7 | 7 | 6 | 6 | 6 | 5 | [Max. DP] |
| **Raffinose** | 3 | 5 | 6 | 6 | 6 | 7 | 7 | 7 | 7 | 7 | 7 | [Max. DP] |

Similar observations were made in the reactions containing raffinose (Table 2, Figure 2 B; Suppl. Figure 1). However, there was no increase in raffinose concentration at extended reaction times, indicating that raffinose-based transglycosylation products are less prone to secondary hydrolysis, as also reflected in the MALDI-TOF results (Table 2). However, MALDI-TOF showed that products of DP 7 only occur later (Table 2), which may be related to the size and structure of raffinose compared to melibiose, hence leaving it a poorer acceptor than melibiose. By reduction of the reducing ends of transglycosylation products the presence of both galactosyl-raffinose oligosaccharides and GOS was confirmed (Suppl. Figure 4), which was expected as raffinose had been shown as a suitable acceptor before (Farkas et al., 2003).

#### 3.1.1. Acceptor-Donor Ratios

Based on the time course experiments, reaction times of 60 min were deemed suitable for all further experiments due to the observed balance between lactose conversion (>62 %), raffinose or melibiose conversion (>21 %) and product size (DP 7). To further favour transglycosylation on melibiose or raffinose instead of lactose, different donor-enzyme (lactose to BgaD-D) and acceptor-donor (melibiose or raffinose to lactose) ratios were tested (Figure 3). Experiments with lactose as the only substrate showed that increased enzyme loading increased the conversion of lactose over the course of an hour (Figure 3 A, B), but the absolute amount of transglycosylation (ΔGlc-Gal) stayed relatively similar while the share of hydrolysis increased (Figure 3 C, D). Addition of melibiose generally increased the total amount of transglycosylation (Figure 3 C) and the conversion of lactose (Figure 3 A), and like in the situation with only lactose present, increased enzyme loading also increased the share of hydrolysis. In all instances, 0.25 µM of BgaD-D showed the longest oligosaccharides (mostly DP 7) and increasing enzyme loading reduced maximum oligosaccharide length by at least 1 DP (Figure 3 C). The same was observed for raffinose (Figure 3 D). Reduced amount of transglycosylation products at increased enzyme loads have also been described for transsialylation (Perna et al., 2021), transfucosylation (Shi et al., 2020) and transglycosylation of N-acetyl-D-glucosamine including *N. circulans* β-galactosidase (Bridiau et al., 2010; Liu et al., 2021). When raffinose was in surplus, product length was also limited to DP 6, most likely due to the higher concentration of acceptors available in solution increasing the chances of interaction with unmodified molecules rather than those already transgalactosylated. Lactose conversion seemed largely unaffected by the addition of raffinose and was only notably increased when raffinose was in excess. However, addition of raffinose did affect the ratio between hydrolysis and transglycosylation: At 0.25 µM BgaD-D, about 70 % of converted lactose led to transglycosylation (based on monosaccharides), but in presence of raffinose, this increased to over 75 %, 85 % or even presumably 100 % (below quantification limits) for ratios 0.25:1, 1:1 and 2:1 of raffinose, respectively (Fig. 2 D). It seems that the share of transglycosylation to melibiose is generally higher than to raffinose, which corroborates the previous observation that melibiose is a better acceptor (Figure 2 A, Table 2). This would also explain why raffinose reactions seemed to profit from excess of acceptor, whilst melibiose worked similar at equimolar concentrations of acceptor and donor. In practice, this would mean that high loadings of raffinose would be desired for transglycosylation efficiency. While raffinose solubility is limited to 203 g/L (0.4 mol/L) at 25.5°C (Hungerford & Nees, 2002; NCBI, 2025b), it increases to 869 g/L (1.7 mol/L) at 50°C (Hungerford & Nees, 2002) . Lactose has a similar solubility at room temperature, 195 g/L (0.5 mol/L, at 20°C) (NCBI, 2025a; Wong & Hartel, 2014), but a concentration of 1 mol/L at 40°C has been used (Warmerdam et al., 2013), showing that concentrations can be increased at elevated temperatures and for scale-up. Melibiose solubility is not a limiting factor as solubilities up to 2500 g/L, equalling > 7 mol/L, (temperature not specified) have been reported (Lakio et al., 2013). In the commercial Vivinal® GOS-preparation made from lactose with the BgaD isomeric mixture, DP 3 accounted for almost a quarter of the whole product mix, while DP 4 was only 10 % (van Leeuwen et al., 2014). When comparing this to results obtained here (Figure 2 C), comparable distributions of oligomers seem to be achieved, although as indicated in Figure 3 (Panel C & D), for some conditions a notable share of transglycosylations is presumably more than single galactosylations (green bars). Apart from the share of different length oligosaccharides, van Leeuwen et al. (2014) also stated that the commercial Vivinal® GOS mixture contains 57% of GOS, 21% of lactose and 22% of monosaccharides. This demonstrates that complete lactose or raffinose conversion is not a requirement for marketing a prebiotic product. In fact, the reactions performed in our study decreased lactose content notably, thus creating a product close to the commercial one. However, for an increased utilization of both lactose and RFOs to reach a higher percentage of galactosyl-RFOs it is worth considering technical solutions like enzyme immobilization or membrane reactors before scaling up. As shown by (Warmerdam et al., 2014), immobilized enzyme in a packed bed reactor can increase productivity of BgaD in GOS production. While (Nath et al., 2013) demonstrated a procedure with immobilization to an ultrafiltration (UF) membrane, using a nanofiltration (NF) membrane of the same material instead could allow collection of shorter oligosaccharides (as small as DP 2-3) as well, whereas UF retains only the enzyme and carbohydrates > 1 kDa MW (DP7, DP6 only with loss) limiting the recovery of the transglycosylated products. Retaining shorter oligosaccharides would also increase repeated interaction of raffinose with BgaD-D and thus a higher share of galactosyl-raffinose products. A possible downside of this is monosaccharide loss, but these could be recovered and used directly or concentrated (after reverse osmosis or drying) as a carbon source for fermentation processes and, in addition, reduces the chance of product inhibition (Boon et al., 1999).

**Figure 3:**
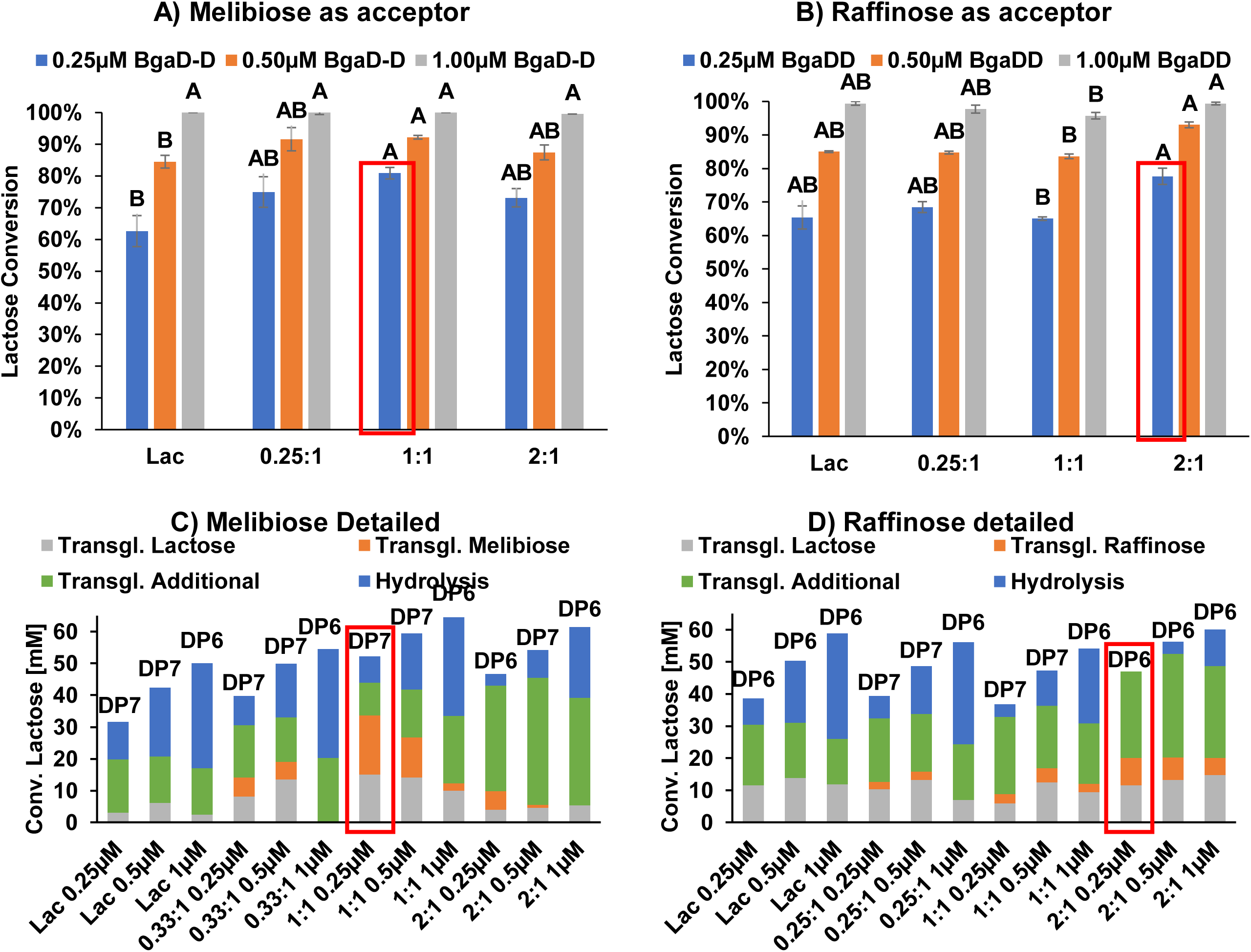
Relative amount of converted lactose as measured by HPAEC-PAD at different additions of **A)** melibiose and **B)** raffinose, respectively: Lac= Lactose only, Ratios 2:1, 1:1, 0.25:1 indicate the amount of melibiose/raffinose compared to Lactose. Reactions were performed with 0.25, 0.50 and 1.00 µM BgaD-D. Error bars denote standard deviations of *n*=3 samples. Upper case letters denote statistically significant differences (*p*<0.05) between samples of the same enzyme loading. **C & D)** The amount of hydrolysis [= galactose release] (blue), transglycosylated melibiose & raffinose [= melibiose & raffinose reduction] (orange), transglycosylated lactose [= converted lactose – glucose release] (grey) and additional transglycosylation to any acceptor [= converted lactose – all previous] (green) as fraction of lactose conversion measured by HPAEC. Labels on top of the bar denote maximum detected oligosaccharide in MALDI-TOF. Reaction conditions: 50°C, 60 min. Red boxes mark reactions with the highest tranglycosylation towards melibiose and raffinose.

### 3.2. Structural characterization by NMR

For structural analysis, reactions were performed at equal concentrations of acceptor and donor (50 mM) and 0.25 µM of BgaD-D (5 µmol BgaD-D / 1 mol lactose) at 50°C. For verbascose, the substrate concentrations were lowered to 10 mM due to its limited availability and enzyme loadings were decreased accordingly. Additionally to pure substrates, an RFO extract from peas was investigated alongside a further purified extract (ethanol precipitated) and one treated with invertase. Products from the reaction with lactose and BgaD-D, as well as transglycosylation products with raffinose and stachyose were separated by preparative HILIC to simplify the NMR-spectra as both HPAEC and MALDI-TOF revealed a multitude of reaction products that would complicate data-interpretation from NMR.

Two oligosaccharides with DP 4 (28 mg, Figure 4 B), which derived from reactions with raffinose as acceptor, could be isolated in quantities which allowed characterization by NMR (Figure 4 A). Furthermore, a fraction of oligosaccharides with DP 5 (4 mg) was obtained. The presence of additional β-galactosides was indicated by doublets around 4.60 ppm with *J*^3^ coupling constants of ∼8 Hz (Karplus, 1963). For one of the isomers with DP 4 (Suppl. Table 1), ^1^H-^13^C-HMBC revealed that transglycosylation had occurred at O-4 of the α-galactose present in raffinose. This substitution at O-4 also caused H-4 and H-1 of said α-galactose to resonate further downfield compared to raffinose (+0.37 ppm for H-4, +0.02 ppm for H-1). For the other isomer with DP 4 (Suppl. Table 2), ^1^H-^13^C-HMBC revealed that transglycosylation had occurred at O-2 of the α-glucose, giving a branched oligosaccharide. This substitution at O-2 also caused the anomeric proton H-1 of α-glucose to resonate 0.16 ppm further downfield. While Farkas et al. (2003) also reported transglycosylation at α-glucose using an unspecified β-galactosidase from *N. circulans*, the preparation used in their study only gave the oligosaccharide which was substituted at O-4 of α-glucose.

**Figure 4.**
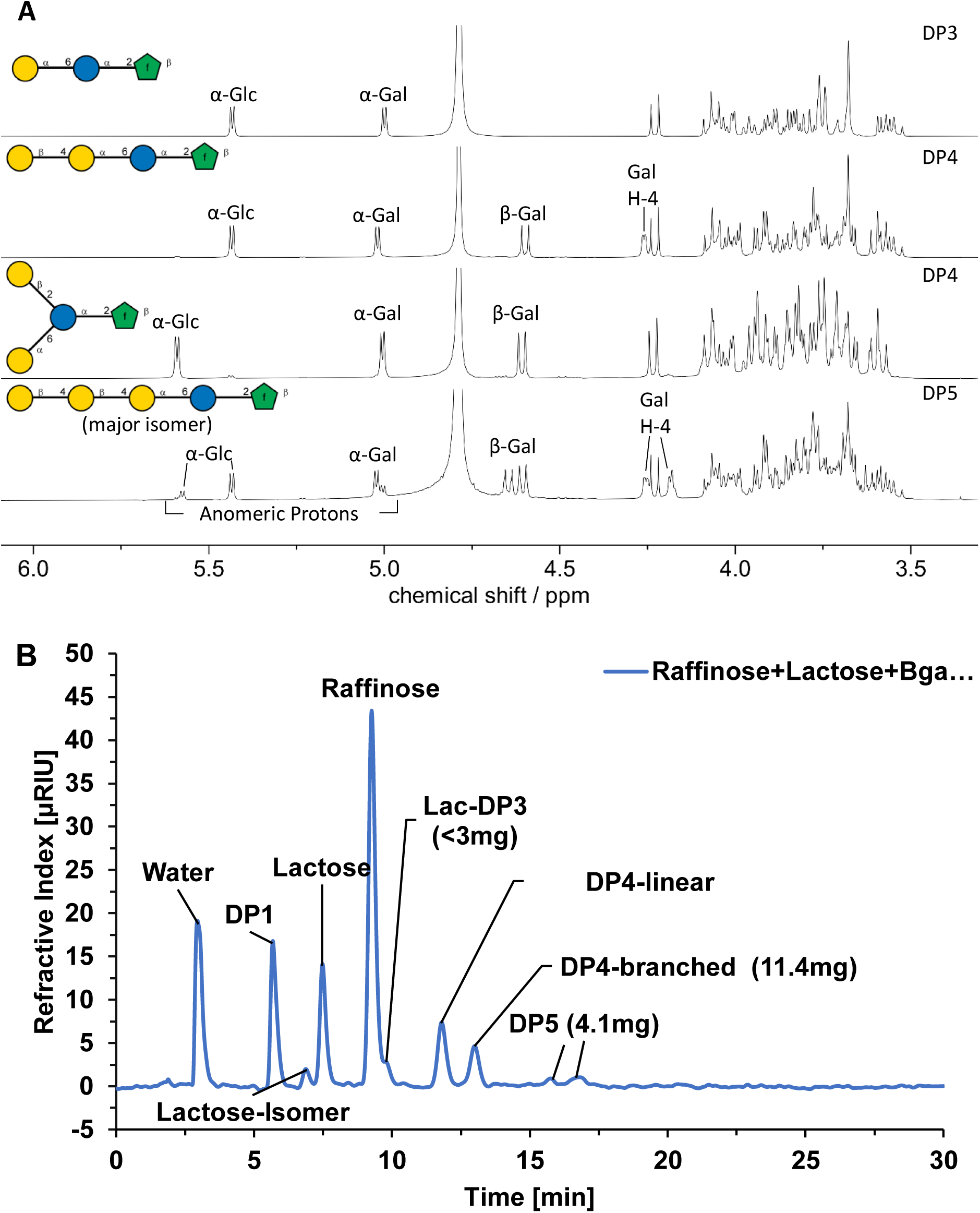
A: ^1^H-NMR (D_2_O, 400 MHz, 298 K) spectra of raffinose (top) and raffinose derived products. **B** Preparative HILIC chromatogram from the purification of raffinose products shown in panel A with annotations of compounds as identified by NMR and obtained weights of these fractions. Lac = Lactose, DP = Monosaccharides.

The differences in product formation could be related to differences in enzyme preparations. According to the CAZy-database (Drula et al., 2022), between 2 and 4 different GH2 enzymes have been found across different *N. circulans* strains. The fraction of oligosaccharides with DP 5 contained a major linear product of which all ^1^H and ^13^C-NMR shifts could be assigned (Suppl. Table 3). The smaller doublets at 5.58 and 5.00 ppm indicate that another branched raffinose derived transglycosylation product with DP 5 is also present. However, the amount of this product was too low to characterize it. For the characterization of oligosaccharides derived from stachyose as acceptor, two fractions were obtained (Figure 5). Like for the products in the raffinose-reactions, the presence of additional galactosides was indicated by doublets between 4.60 and 4.70 ppm with *J*^3^ coupling constants of ∼8 Hz. The fraction containing oligosaccharides with DP = 5 (31.2 mg) consisted of a 3:2 mixture of two isomers. Despite the complexity of the mixture, a complete assignment of all ^1^H and ^13^C-NMR shifts for both isomers was possible (Suppl. Table 4-5). For both isomers, transglycosylation exclusively occurred at O-4 of either of the two α-galactoses present in stachyose resulting in the formation of a mixture of linear and branched pentasaccharides (Figure 5 A). In contrast to the transglycosylation products derived of raffinose, no galactose substituents were observed at O-2 of the glucose. The fraction containing oligosaccharides with DP = 6 (6.9 mg) was too complex for complete assignment of all ^1^H and ^13^C-NMR shifts. However, HSQC-TOCSY showed the presence of both O-4 substituted and non-O-4 substituted α-galactoses and revealed the presence of four non-O-4 substituted and two O-4 substituted β-galactoses (Figure 6 A). In ^1^H-^13^C-HMBC correlations from the non-substituted β-galactoses to O-4 of both α-galactoses and β-galactoses could be observed suggesting that the sample contained a mixture of three different oligosaccharides shown in Figure 6 B.

**Figure 5.**
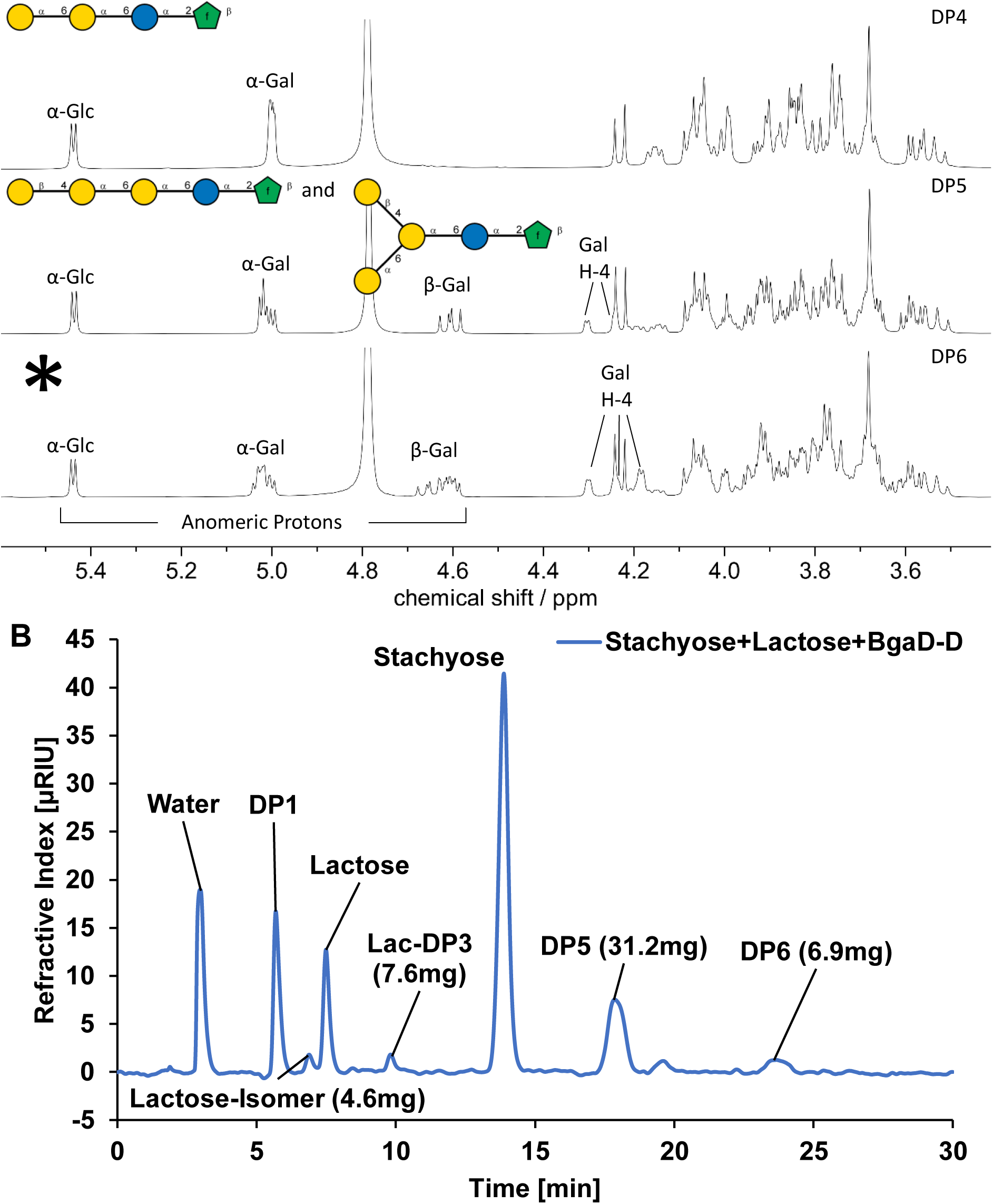
A: ^1^H-NMR (D_2_O, 400 MHz, 298 K) spectra of stachyose (top) and stachyose derived products. *=Structure of DP6 elucidated in Figure 6. **B:** Preparative HILIC chromatogram from the purification of stachyose products shown in panel A with annotations of compounds as identified by NMR and obtained weights of these fractions. Lac = Lactose, DP1 = Monosaccharides.

**Figure 6.**
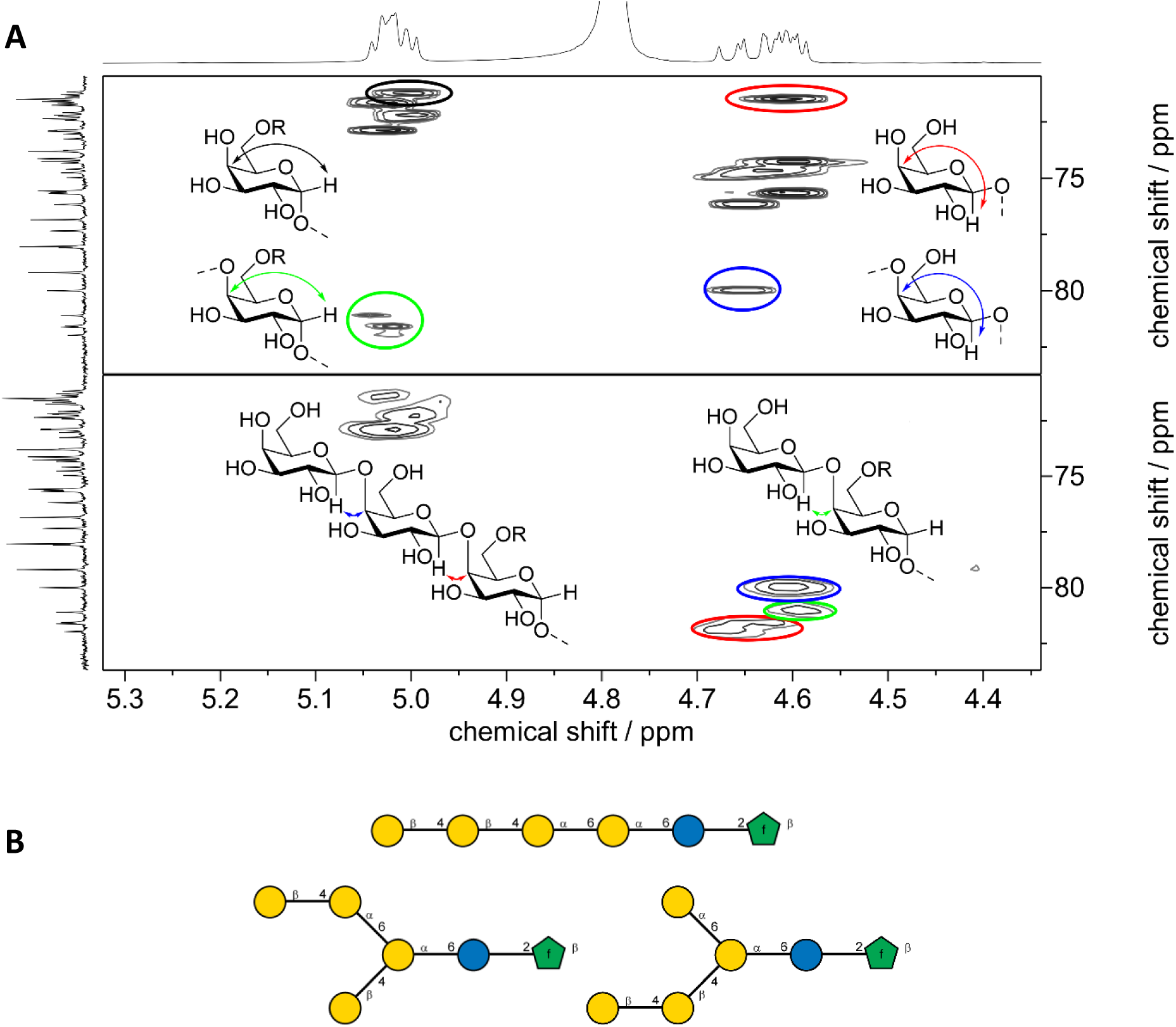
A: HSQC-TOCSY (top) showing the presence of both terminal and internal α- and β-galactoses in stachyose derived products with DP 6. HMBC (bottom) showing correlations of β-galactoses to O-4 of α- and β-galactoses. **B:** proposed structures of stachyose derived products with DP 6.

From the reaction involving just lactose and BgaD-D 1-4 β-linked oligomers with up to DP 4 could be isolated in quantities which allowed characterization by NMR (Suppl. Figure 5). The NMR data of these oligomers in (Suppl. Table 6-8) agreed with the data reported in a previous study (van Leeuwen et al., 2014), confirming their identity. Both, the tri- and tetrasaccharide (Suppl. Figure 5 A) have been reported as major BgaD-D products before (Yin et al., 2017a). A regioisomer of lactose bearing a β-1,3 linkage, which also has been described in an earlier study by Yin et al. (2017a), was also identified. Both the lactose-isomer and DP3 from lactose were also found in reactions with raffinose and stachyose, confirming that lactose also works as acceptor in these reactions, although to a lesser extent than the RFOs. It is worth highlighting that the fractions of mixed-linkage oligosaccharides from raffinose (Figure 4 B) and stachyose (Figure 5 B) were clearly larger than those of lactose-derived products (by absolute weight), indicating that these reactions lead to a notable share of galactosylated RFO’s, as also indicated by the decrease of RFOs in time-course experiments (Figure 1). Furthermore, when using melibiose and complex mixtures of RFO substrates (RFO-extract, invertase treated RFO-extract and ethanol precipitated RFO-extract), the same indicators for substitution on the α-galactoses and the O-4 position were detected by NMR (Supplementary Figure 6) as for raffinose and stachyose, strongly suggesting that all RFO substrates are acting as acceptors in reactions with lactose and BgaD-D, also in mixtures.

### 3.3. Microbial fermentations

We assessed the fermentability of the novel RFO-derived oligosaccharides by a variety of food and gut-related microorganisms (eleven strains in total) to evaluate any potential prebiotic effects (Figure 7, Figure 9; Supplementary Figure 7).

**Figure 7:**
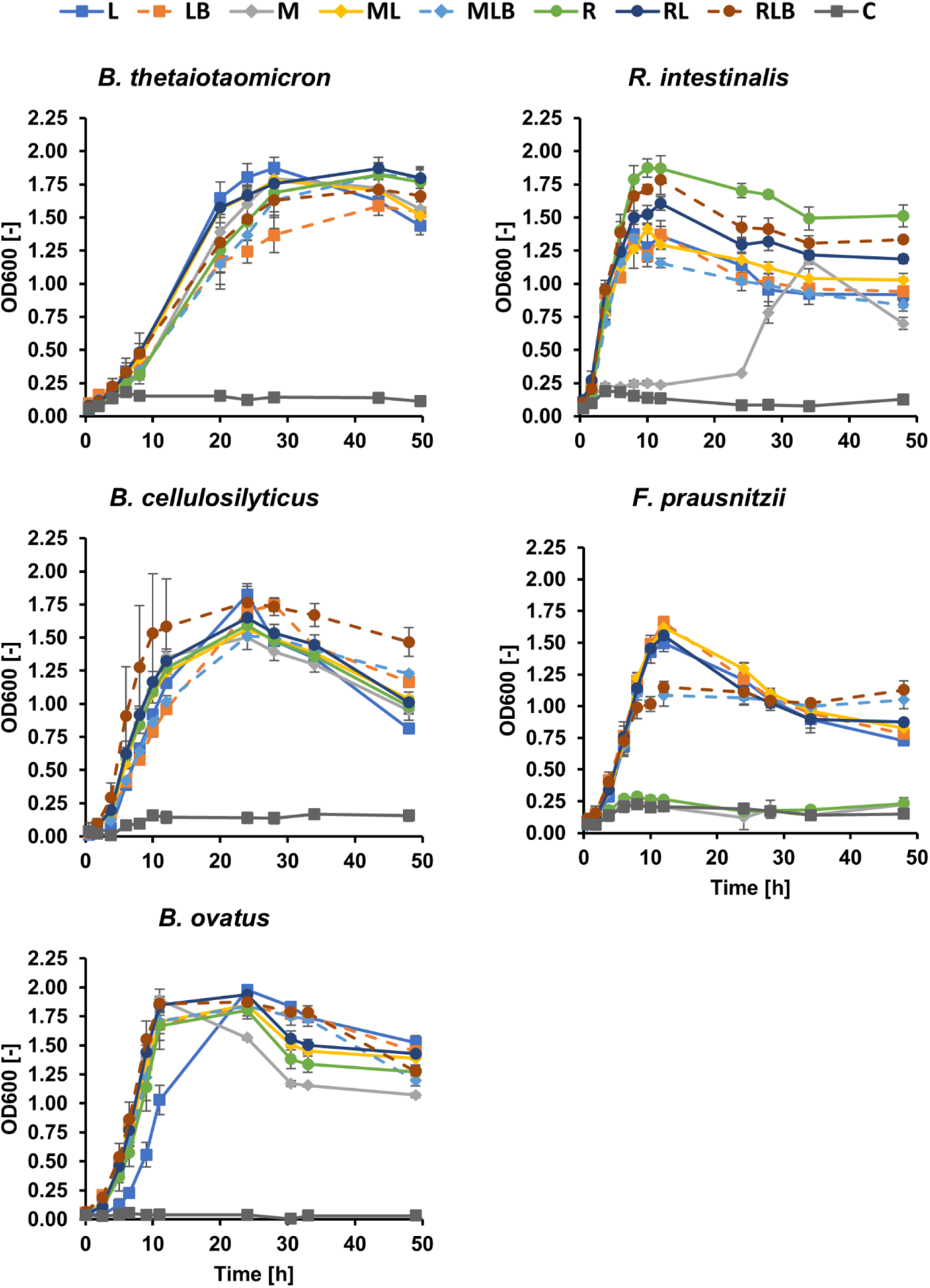
Average growth curves (*n*=3) of all bacterial strains fermented with different carbon sources at anaerobic conditions and 37°C. Error bars denote standard deviation for n=3 samples. L = Lactose, LB = Lactose + BgaD-D, M = Melibiose, ML = Melibiose + Lactose, MLB = Melibiose+ Lactose+BgaD-D, R=Raffinose, RL = Raffinose+Lactose, RLB = Raffinose+Lactose+BgaD-D.

#### 3.3.1. Fermentations on unpurified transglycosylation products

Due to a simpler workup than HILIC purified oligos, and thus allowing larger production, fermentation on unpurified mixtures obtained after transglycosylation was done as a first screening for all strains in this study. Relevant gut bacteria were used in anaerobic fermentations (Figure 7) and formed two clear groups of strains. The first group consists of three *Bacteroides* spp. strains, which grew well on all substrates and reached similar maximum culture densities (OD_600_ > 1.8) within 24 hours and was accompanied by a pH decline from 7 initially to approximately 6 after 48 hours. Controls without a carbon source had minimal growth (OD_600_ < 0.2) and no pH decline.

*Bacteroides thetaiotaomicron* had the longest lag-phase but reached comparable cell densities after 24 hours. Some cultures had detectable carbohydrate residues remaining after the fermentations (as analyzed by HPAEC, Figure 8 A): *Bacteroides ovatus* and *Bacteroides cellulosilyticus* both had remaining monosaccharides (the latter strain at low concentrations), and while *B. thetaiotaomicron* utilized the majority of raffinose (approx. 90 %), only residues of melibiose were detected in solution, indicating both invertase & α-galactosidase activity (GH32 & GH36 present in the genome). In some of the BgaD-D treated solutions residues of lactose, melibiose or their potentially galactosylated derivatives were observed (Figure 8 A), and masses corresponding to tri- and tetrasaccharides (527 / 543 and 689 / 705 *m/z*, respectively) were detected in MALDI-TOF MS after cultivation of *B. thetaiotaomicron* (Figure 8 B), corresponding to the slightly reduced growth on more complex substrates (Figure 7). The second group of strains consisted of *Roseburia intestinalis* and *Faecalibacterium prausnitzii*, as these strains showed either no or only delayed growth on some of the mixed oligosaccharides (LB, MLB, RLB). In samples with no growth, pH remained the same as in pure media (pH 7 for YCFA & Minimal media). *R. intestinalis* clearly grew best on raffinose-containing carbon sources, (OD_600_ 1.6-1.8), whereas lactose-containing media had slightly lower growth (OD_600_ 1.3-1.4). Pure melibiose was utilized with a delay of 24 hours (Figure 7). For *R. intestinalis* grown on LB some trisaccharide residues were detected in MALDI-TOF spectra and the corresponding HPAEC chromatograms, whereas this could not be observed for RFO-derived samples (MLB, RLB) did not.

**Figure 8.**
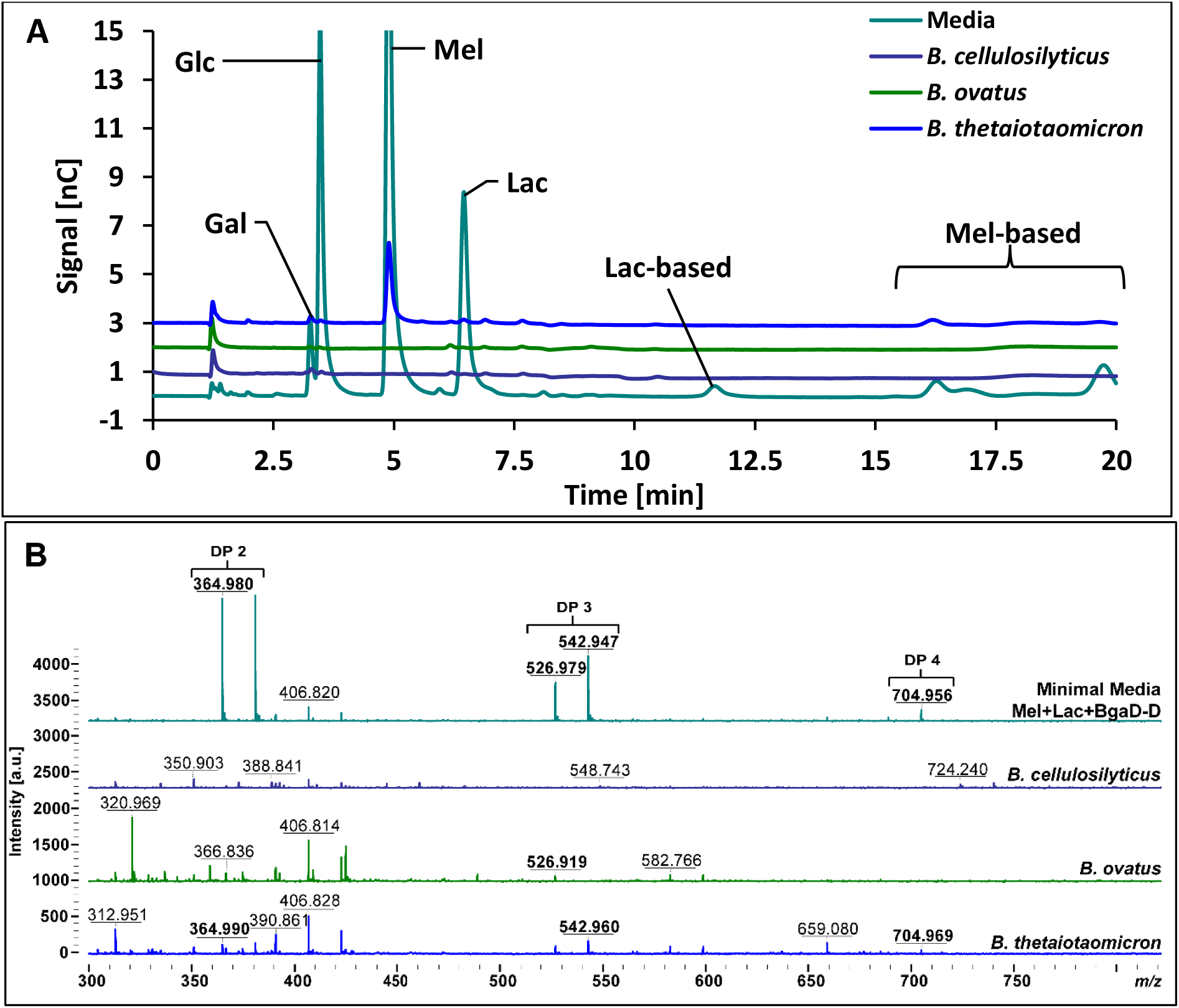
A: HPAEC-PAD chromatograms for the same samples as in panel A with annotations of known peaks. Symbols: Gal=Galactose, Glc=Glucose, Mel=Melibiose, Lac=Lactose, Lac-base=Lactose transglycosylation product, Mel-based=Melibiose transglycosylation product. **B:** MALDI-TOF spectra of all *Bacteroides* strains in media supplemented with BgaD-D treated melibiose+lactose (MLB) after 48 hours with oligosaccharides highlighted in bold and

**Figure 9:**
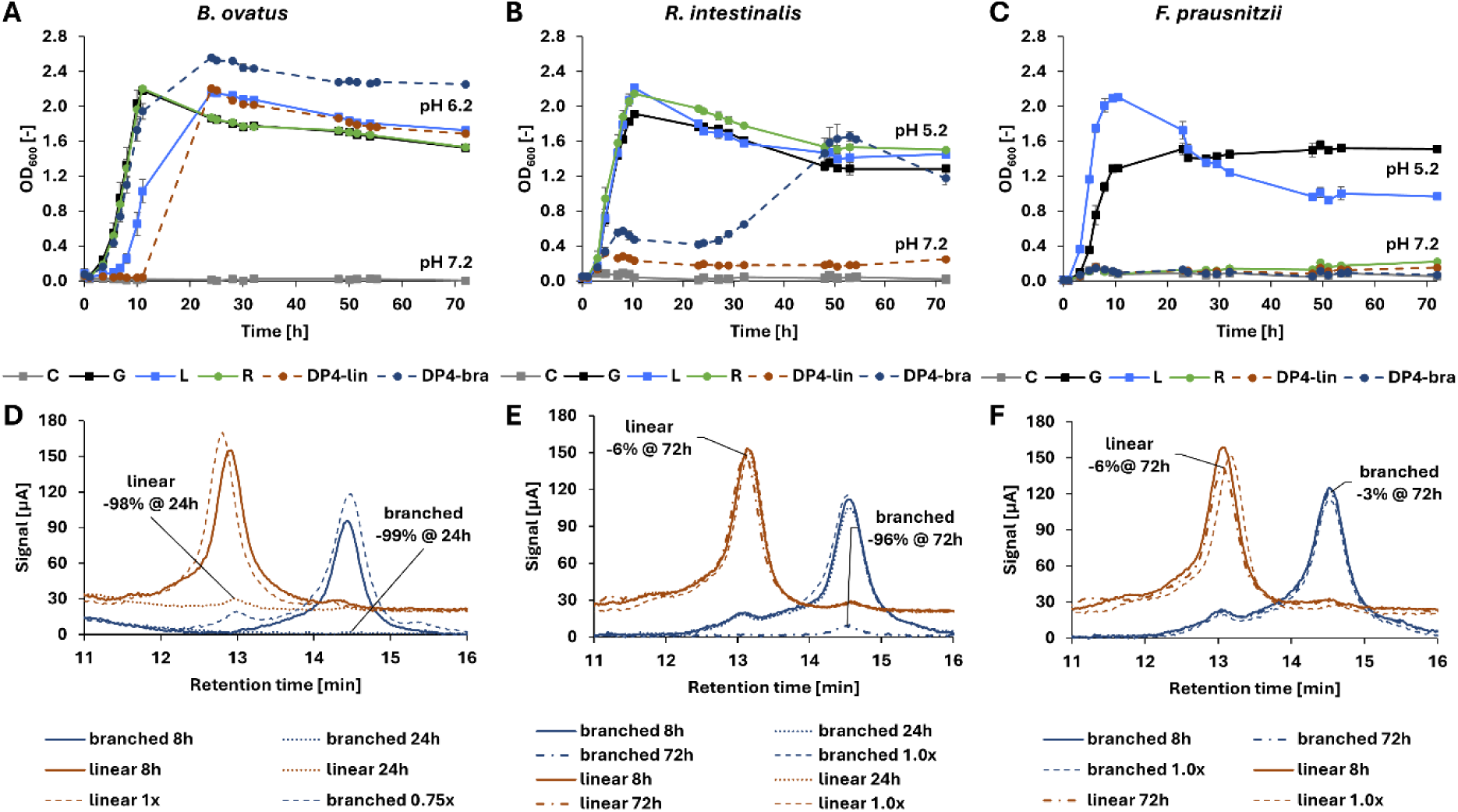
Results from fermentations with purified linear and branched galactosyl-raffinose (tetrasaccharides; Figure 4) in combination with *Bacteroides ovatus*, *Roseburia intestinalis*, and *Fecalibacterium prausnitzii*. **Panels A-C** show the respective growth curves over 72 hours (*n*=3 with standard deviation) in comparison to glucose, lactose, raffinose and no carbon source. The pH at the final time point is indicated. **Panels D-F** show a zoom of the analytical HILIC chromatograms for the tetrasaccharides at relevant time points (*n=1*) for all three strains, respectively. Linear tetramer chromatograms are offset with +20µA for visibility. Dashed lines represent unfermented references. The percentual reductions are based on triplicates and a 5-point calibration curve.

For *F. prausnitzii*, the trend was almost inverse to *R. intestinalis*: cultures only containing raffinose and melibiose had negligible growth compared to the control. Lack of growth on raffinose may be explained by the fact that *F. prausnitzii* is missing the necessary invertase (GH32, Table 1). With lactose in the media, it did, however, grow well (OD_600_ = 1.6), and pH was reduced (pH 4-5). Interestingly, BgaD-D treatment of RFOs (MLB, RLB) reduced growth (OD_600_ = 1.1), but not for lactose + BgaDD (LB, OD_600_ = 1.7). One may hypothesize that *F. prausnitzii* does not take up galactosylated melibiose and raffinose (or pure melibiose/raffinose for that matter), even though their β-galactosidase could hydrolyze it (see L, LB Figure 7). Indeed, these galactosidases are usually not secreted and only active on the substrate upon uptake into the cell, as was shown for the degradation of mannan in which *F. prausnitzii* is reliant on other organisms secreted enzymes (Lindstad et al., 2021). Unchanged melibiose and raffinose concentrations were confirmed by HPAEC-PAD. All strains that anaerobically utilized raffinose and/or melibiose (*Bacteroides spp.* & *R. intestinalis*) have GH27 and GH36 α-galactosidases in their genome (Table 1), and the two families contain structurally similar enzymes (Reddy et al., 2016). It has been suggested that some of the GH36s in strains of *B. ovatus* (Reddy et al., 2016), *R.* intestinalis (La Rosa et al., 2019) and in *F. prausnitzii* (Lindstad et al., 2021) are related to polysaccharide utilization loci for mannan-degradation and are active on galactose side-chains of galactomannans. It is therefore possible that some of these GH36s are not transcribed unless mannan is present to induce the relevant activities. Thus, they might not be active on raffinose or melibiose alone. Additionally, these enzymes are intracellular (La Rosa et al., 2019; Lindstad et al., 2021) and non-fermenters (*F. prausnitzii*) might be missing a relevant transporter. The *B. thetaiotaomicron* strain used here additionally harbours 11 annotated GH97s which could add to its capability to degrade RFOs, as one of the *B. thetaiotaomicron* GH97 has been reported to be active on melibiose and raffinose (Shin et al., 2020). The *B. ovatus* strain has 12 different GH97s in its genome (Drula et al., 2022) and grew notably in these experiments (Figure 7). Furthermore, it is worth noting that according to the recent division of family GH2 (van Zanten et al., 2012) BgaD-D is in the same subfamily (#1) as characterized enzymes of *B. ovatus* and *B. thetaioaomicron*, supporting the observation that *Bacteroides* spp. can utilize the produced α-β-GOS.

Investigating aerobic fermentations are also relevant because food microorganisms and administered probiotics are generally added in environments that are either aerobic or at least not strictly anaerobic, meaning that oxygen-tolerant probiotic organisms could potentially utilize the produced oligosaccharides in food products and before reaching the consumer’s gut.

Like the anaerobic cultivations, these aerobic fermentations with food-related strains showed no unified response to BgaD-D-treated substrates. Nevertheless, two groups emerge from the results: *Levilactobacillus brevis and Lactiplantibacillus pentosus* KW1 and KW2 were among the best growing strains (OD_600_ = 1.4-2.1) compared to controls (Suppl. Figure 7) but did prefer different carbon sources. *L. brevis* cultures grown on untreated carbon sources (L, R, M, ML, RL) reached slightly higher densities then their BgaD-D treated counterparts (ΔOD_600_=0.1-0.3). For lactose (L, LB), this difference disappeared towards 48 hours. The delayed and reduced growth on some substrates might be due to the necessity for multiple enzymes (additional β-linkage) or reduced affinities of modified oligosaccharides to the degrading enzymes and/or respective transporters (e.g., L/ML/RL compared to LB/MLB/RLB). However, that *L. brevis* grew well on all substrates supports previous findings that some lactobacilli strains have α-galactosidases, potential RFO transporter proteins and regulators as well as β-galactosidases in close proximity in their genome, if not in the same locus (Hachem et al., 2012). For *L. pentsosus* KW1, a slightly different picture arose: Cultures grew better on BgaD-D treated substrates (OD_600_ = 1.3-1.6) than their untreated counterparts (OD_600_ = 1.1-1.3), with the biggest difference for lactose (L vs. LB). This indicates either better monosaccharide uptake and/or a better suited and readily available β-galactosidase than in *L. brevis*. The general utilization of all substrates for growth by *L. pentsosus* KW1 is in line with the presence of all relevant enzymes in the genome to degrade RFOs and lactose (Lombard et al., 2014; Wiull et al., 2024) (Table 1). Interestingly, *L. pentosus* KW2 was isolated from the same environment as *L. pentosus* KW1 (Wiull et al., 2024) and has the same enzymatic toolbox (Table 1), but showed different substrate preferences (Suppl. Figure 7): Melibiose-containing media stimulated most growth (M, ML, MLB) whereas raffinose seemed to be the least preferred substrate as cultures grew like the control. Growth on RFOs was expected for all three strains as they showed good response in a previous study with RFO extract from peas (Garbers et al., 2025). The results in our study, however, add some detail and show different preferences between melibiose and raffinose for the *L. pentosus* strains. The second group, consisting of *Lacticaseibacillus rhamnosus* GG (also ATCC 53103) and *Lactococcus lactis subsp. cremoris*, displayed overall notably lower growth in controls (OD_600_ = 0.5-0.6) and with carbohydrates (OD_600_ = 0.7-0.9). The only carbon sources leading to higher final densities than controls for both strains were LB and RL. Both strains also showed comparably slow growth and only little benefit of RFO additions in a previous study (Garbers et al., 2025). *L. rhamnosus* GG was isolated from human faeces (Segers & Lebeer, 2014) and is a commonly applied probiotic in food and medicine (Banna et al., 2017). Despite the strain having genetic prerequisites for degrading lactose and RFOs (Table 1), Huang et al. (2024) showed that RFOs from chickpea aquafaba did not stimulate growth of *L. rhamnosus*. This could be considered an advantage, as raffinose-derived prebiotics could be used in conjunction with this strain to both supplement a consumer with pro- and prebiotics without risking fermentation during storage. Lastly, *Lentilactobacillus buchneri* appears to be an intermediate between these two groups. Control cultures grew like the first group (OD_600_ ≥ 1.0), but additional carbon sources only stimulated growth after > 40 hours and all grew to a similar extend (approx. OD_600_ = 1.2) apart from RL (OD_600_ > 1.4) and L (OD_600_ < 0.8). Increased growth in presence of RFOs was previously shown (Garbers et al., 2025), but the low response to lactose (L) is surprising since the strain has 3 annotated GH2 enzymes (β-galactosidase) in its genome (Table 1) and it grew well on LB. However, compared to *L.* pentosus and *L.* brevis, it does not harbour any GH42 and since the strain is from silage (Dysvik et al., 2020; Heinl et al., 2012), it might have adapted to other carbon sources, whereas lactose is uncommon in these environments. Carbohydrate analysis of the aerobic samples proved to be non-trivial due to notable noise from media (Suppl. Figure 8).

Taken together, there is no clear trend that certain oligosaccharides are generally indigestible, but rather that certain strains have preferences of different substrates, except for the *Bacteroides* strains that grew comparably well on all substrates. Indeed, some *Bacteroides* strains have some of the biggest toolboxes for carbohydrate degradation (Shin et al., 2020), and in this case it appeared that the number of relevant CAZymes can be an indicator of degradation potential (Table 1). It seems, however, that adding an additional linkage to the oligosaccharides and increasing their length might create some hurdles for either uptake or enzymatic degradation for some of the bacteria as indicated by the differences in growth.

#### 3.3.2. Fermentations on purified transglycosylation products

Realizing that potential background growth of mono- and disaccharides caused difficulties in interpreting fermentability, we decided to purify a larger batch of the two tetramers from raffinose (Figure 4) to get a more detailed picture on the fermentability of these. Purification of the two major products from raffinose-based reactions (galactosyl-raffinose tetramers, Figure 4) was conducted with preparative scale HILIC. To obtain approx. 200 mg of each tetramer, a reaction with 5 g substrates (or 115 mL reaction volume) was prepared and purified via 20 injections on the preparative HILIC. Based on the results obtained with the product mixtures above, we chose three organisms for anaerobic fermentations with the purified tetramers, specifically; *B. ovatus* (generally grew well), *R. intestinalis* (raffinose utilizer), and *F. prausnitzii* (non-raffinose utilizer). Lactose and raffinose were included for comparison. Results confirm the previous conclusions that *B. ovatus* metabolizes all substrates for growth (OD ^max^>2 and >98% reduction in 24 hours) (Figure 9). Notably, there is a delay for the utilization of the linear tetramer (-5% at 8 h) compared to the branched tetramer (-32 % at 8 h) which might indicate a lower availability for the readily expressed hydrolases and active systems as discussed previously. For *R. intestinalis*, both modified substrates initially posed a challenge, and growth was notably reduced, but larger than in the control culture and the branched-DP4 had a 13 % decrease at 8h and even 21% decrease at 24 hours according to HILIC. From 27 hours onwards the strain also increasingly grew to comparable levels as the other substrates and at 48 h 63 % of the substrates were consumed, ultimately reaching 96 % after 72 h. The linear tetramer was barely utilized by *R. intestinalis* (OD_600_^max^ = 0.28), and concentrations decreased only by 6 %. The delay in growth of *B. ovatus* and the low response of *R. intestinalis* to the linear tetramer might indicate that GH32s (Invertase) are not as readily active as α-galactosidase (GH36) in these strains, as GH36 is inactive on the linear tetramer, while GH32 is inactive on the branched tetramer (Supplementary Figure 9, 10). The activity of GH36s is also indicated by the clear growth on raffinose (no residues detected in either strain). β-galactosidases (e.g., GH2) can also still play a role as at least BgaD-D is able to hydrolyse either of the products (Supplementary Figure 11). As observed before, *F. prausnitzii* did not grow on raffinose and neither on its derivatives as indicated by the clear reduction in the observed maximum OD_600_ (cf. Fig. 6 and Fig. 8). Analysis confirmed that all three substrates were indeed present at concentrations close to the starting point at 72 hours; for the linear and branched tetramer reductions of only 6 and 3 % were measured, respectively. This demonstrates that the differences in growth observed during the first screening (Fig. 6) can indeed be linked to the reduced fermentability of some of the transglycosylation products. While the environment in the gut will be largely different to the here performed *in vitro* fermentations, these tests nevertheless demonstrate that some of the tested commensal gut bacteria are able to utilize the raffinose derivatives whilst others are either not able to metabolize them or at least show a delayed response due to the uncommon nature of the carbohydrate. This could result in a reduction of the rapid fermentation response as described by the FODMAP-hypothesis (Gibson & Shepherd, 2005) and increases specificity of these carbohydrates which is a key requirement for prebiotics (Gibson et al., 2017). This is strengthened by previous studies hinting at especially (α1-6), (β1-2) and (β1-4), all present in the tested α-β-GOS, to show prebiotic potential in disaccharides (Onyango et al., 2023; Sanz et al., 2005).

## 4. Conclusion

In this study we showed that RFOs work as efficient acceptors for transglycosylation with lactose and BgaD-D leading to new mixed-linkage α-β-GOS. RFOs with additional galactose (β1-4) linked to the terminal galactose were the predominant products of reactions. The transglycosylation activity increased when enzyme loadings were kept low (e.g., 5 µmol BgaD-D / 1 mol lactose) and acceptors other than lactose were available at equal concentrations (melibiose) or in excess (raffinose).

Fermentations with BgaD-D treated substrates indicated that some commensal gut bacteria like *Bacteroides* spp. easily utilize the α-β-GOS presented herein and grow notably; for *B. ovatus* full utilization of two purified α-β-GOS could be demonstrated. Other strains, including those with smaller enzymatic toolboxes, have reduced growth and face some challenges in utilizing the oligosaccharides; for *F. prausnitzii* it could be shown that neither of the purified α-β-GOS could be fermented. These results might imply a reduced FODMAP potential while simultaneously showing that the products are not completely indigestible.

Although the here performed *in vitro* fermentations of pure strains give valuable information, additional studies with either faecal samples or model systems of the digestive tract could help to better understand the effect of the produced oligosaccharides on consumers compared to pure raffinose or lactose and well-known prebiotics. Furthermore, scale-up and further optimization of reaction conditions (e.g., immobilization or use of membrane reactors) could give an insight into the industrial feasibility of the process, especially in comparison to the established GOS production.

## Acknowledgements

The authors would like to thank Jesper Holck (DTU) for support with the carbohydrate analysis facilities, Ingrid Rokke Elvebakken (NMBU) for support with enzyme productions and Catrin Tyl (NMBU) for support with statistical analysis. Further thanks to all colleagues providing bacterial stocks: Sabina Leanti La Rosa, Lars Lindstad, Hilde Østlie, Davide Porcellato, Geir Mathiesen and Kamilla Wiull (all NMBU).

## CRediT authorship contribution statement

**Philipp Garbers:** Conceptualization, Methodology, Investigation, Writing – original draft, review & editing, funding acquisition. **Gordon J. Boehlich:** Methodology, Investigation, Writing – original draft, review & editing. **Birgitte Zeuner:** Conceptualization, Methodology, Supervision, Writing –review & editing; **Jane W. Agger:** Conceptualization, Methodology, Supervision, Writing –review & editing. **Bjørge Westereng:** Conceptualization, Methodology, Supervision, Writing –review & editing, funding acquisition, project administration.

## References

1. BacDrive. 2025. Bacteroides cellulosilyticus DSM 14838.

2. Banna, G.L., Torino, F., Marletta, F., Santagati, M., Salemi, R., Cannarozzo, E., Falzone, L., Ferrau, F., Libra, M. 2017. Lactobacillus rhamnosus GG: An Overview to Explore the Rationale of Its Use in Cancer. Frontiers in Pharmacology, 8, 603.

3. Barile, D., Rastall, R.A. 2013. Human milk and related oligosaccharides as prebiotics. Current Opinon in Biotechnology, 24(2), 214–9.

4. Bhatty, R.S., Christison, G.I. 1984. Composition and nutritional quality of pea (Pisum sativum L.), faba bean (Vicia faba L. spp. minor) and lentil (Lens culinaris Medik.) meals, protein concentrates and isolates. Qualitas Plantarum Plant Foods for Human Nutrition, 34(1), 41–51.

5. Boon, M.A., Janssen, A.E.M., van der Padt, A. 1999. Modelling and parameter estimation of the enzymatic synthesis of oligosaccharides by ?-galactosidase fromBacillus circulans. Biotechnology and Bioengineering, 64(5), 558–567.

6. Bridiau, N., Issaoui, N., Maugard, T. 2010. The effects of organic solvents on the efficiency and regioselectivity of N-acetyl-lactosamine synthesis, using the beta-galactosidase from Bacillus circulans in hydro-organic media. Biotechnolog Progress, 26(5), 1278–89.

7. Daveby, Y.D., Abrahamsson, M., Åman, P. 2006. Changes in chemical composition during development of three different types of peas. Journal of the Science of Food and Agriculture, 63(1), 21–28.

8. Demarco, A., Gibon, V. 2020. Overview of the soybean process in the crushing industry. Ocl, 27.

9. Drula, E., Garron, M.L., Dogan, S., Lombard, V., Henrissat, B., Terrapon, N. 2022. The carbohydrate-active enzyme database: functions and literature. Nucleic Acids Research, 50(D1), D571–D577.

10. Dysvik, A., La Rosa, S.L., Liland, K.H., Myhrer, K.S., Ostlie, H.M., De Rouck, G., Rukke, E.O., Westereng, B., Wicklund, T. 2020. Co-fermentation involving *Saccharomyces cerevisiae* and *Lactobacillus* species tolerant to brewing-related stress factors for controlled and rapid production of sour beer. Frontiers in Microbiology, 11, 279.

11. Elango, D., Rajendran, K., Van der Laan, L., Sebastiar, S., Raigne, J., Thaiparambil, N.A., El Haddad, N., Raja, B., Wang, W., Ferela, A., Chiteri, K.O., Thudi, M., Varshney, R.K., Chopra, S., Singh, A., Singh, A.K. 2022. Raffinose family oligosaccharides: Friend or foe for human and plant health? Frontiers in Plant Science, 13, 829118.

12. Farkas, E., Schmidt, U., Thiem, J., Kowalczyk, J., Kunz, M., Vogel, M. 2003. Regioselective Synthesis of Galactosylated Tri- and Tetrasaccharides by Use of β-Galactosidase from *Bacillus circulans*. Synthesis(5), 0699–0706.

13. Garbers, P., Brandal, H.A., Vardeberg Skeie, A., Karlsnes, G.W., Varela, P., Tyl, C., Westereng, B. 2025. Pea-Derived Raffinose-Family Oligosaccharides as a Novel Ingredient to Accelerate Sour Beer Production. Journal of Agricultural and Food Chemistry, 73(7), 4219–4230.

14. Garbers, P., Gaber, S., Knezevic, D., Tyl, C., Sahlstrøm, S., Knutsen, S., Westereng, B. 2026. A pilot-scale process for the extraction of raffinose-oligosaccharides from pulse protein concentrates. Innovative Food Science & Emerging Technologies, 108.

15. Gibson, G.R., Hutkins, R., Sanders, M.E., Prescott, S.L., Reimer, R.A., Salminen, S.J., Scott, K., Stanton, C., Swanson, K.S., Cani, P.D., Verbeke, K., Reid, G. 2017. Expert consensus document: The International Scientific Association for Probiotics and Prebiotics (ISAPP) consensus statement on the definition and scope of prebiotics. Nature Reviews Gastroenterology & Hepatology, 14(8), 491–502.

16. Gibson, P.R., Shepherd, S.J. 2005. Personal view: food for thought--western lifestyle and susceptibility to Crohn’s disease. The FODMAP hypothesis. Alimentary Pharmacology and Therapeutics, 21(12), 1399–409.

17. Gulewicz, P., Ciesiolka, D., Frias, J., Vidal-Valverde, C., Frejnagel, S., Trojanowska, K., Gulewicz, K. 2000. Simple method of isolation and purification of alpha-galactosides from legumes. Journal of Agricultural and Food Chemistry, 48(8), 3120–3.

18. Gänzle, M.G. 2012. Enzymatic synthesis of galacto-oligosaccharides and other lactose derivatives (hetero-oligosaccharides) from lactose. International Dairy Journal, 22(2), 116–122.

19. Hachem, M.A., Fredslund, F., Andersen, J.M., Jonsgaard Larsen, R., Majumder, A., Ejby, M., Van Zanten, G., Lahtinen, S.J., Barrangou, R., Klaenhammer, T., Jacobsen, S., Coutinho, P.M., Lo Leggio, L., Svensson, B. 2012. Raffinose family oligosaccharide utilisation by probiotic bacteria: insight into substrate recognition, molecular architecture and diversity of GH36 α-galactosidases. Biocatalysis and Biotransformation, 30(3), 316–325.

20. Heinl, S., Wibberg, D., Eikmeyer, F., Szczepanowski, R., Blom, J., Linke, B., Goesmann, A., Grabherr, R., Schwab, H., Puhler, A., Schluter, A. 2012. Insights into the completely annotated genome of Lactobacillus buchneri CD034, a strain isolated from stable grass silage. Journal of Biotechnology, 161(2), 153–66.

21. Hovorkova, M., Kascakova, B., Petraskova, L., Havlickova, P., Novacek, J., Pinkas, D., Gardian, Z., Kren, V., Bojarova, P., Smatanova, I.K. 2024. The variable structural flexibility of the Bacillus circulans beta-galactosidase isoforms determines their unique functionalities. Structure, 32(11), 2023–2037 e5.

22. Huang, Y.P., Masarweh, C., Paviani, B., Mills, D.A., Barile, D. 2024. Exploring bioactive compounds in chickpea and bean aquafaba: Insights from glycomics and peptidomics analyses. Food Chemistry, 460(Pt 2), 140635.

23. Hungerford, E.H., Nees, A.R. 2002. Raffinose - Prepartion and Properties. Industrial & Engineering Chemistry, 26(4), 462–464.

24. Jameson, J.K., Mathiesen, G., Pope, P.B., Westereng, B., La Rosa, S.L. 2021. Biochemical characterization of two cellobiose 2-epimerases and application for efficient production of lactulose and epilactose. Current Research in Biotechnology, 3, 57–64.

25. Karplus, M. 1963. Vicinal Proton Coupling in Nuclear Magnetic Resonance. Journal of the American Chemical Society, 85(18), 2870–2871.

26. Kim, S., Kim, W., Hwang, I.K. 2003. Optimization of the extraction and purification of oligosaccharides from defatted soybean meal. International Journal of Food Science & Technology, 38(3), 337–342.

27. La Rosa, S.L., Leth, M.L., Michalak, L., Hansen, M.E., Pudlo, N.A., Glowacki, R., Pereira, G., Workman, C.T., Arntzen, M.O., Pope, P.B., Martens, E.C., Hachem, M.A., Westereng, B. 2019. The human gut Firmicute Roseburia intestinalis is a primary degrader of dietary beta-mannans. Nature Communications, 10(1), 905.

28. Lakio, S., Sainio, J., Heljo, P., Ervasti, T., Kivikero, N., Juppo, A. 2013. The tableting properties of melibiose monohydrate. International Journal of Pharmaceutics, 456(2), 528–35.

29. Lebreton, A., Garron, M.L., Vuillemin, M., Pilgaard, B., Hornung, B.V.H., Drula, E., Lombard, V., Helbert, W., Henrissat, B., Terrapon, N. 2025. Division of the large and multifunctional glycoside hydrolase family 2: high functional specificity and biochemical assays in the uncharacterized subfamilies. Biotechnology for Biofuels and Bioproducts, 18(1), 68.

30. Leivers, S., Lagos, L., Garbers, P., La Rosa, S.L., Westereng, B. 2022. Technical pipeline for screening microbial communities as a function of substrate specificity through fluorescent labelling. Nature Communications Biology, 5(1), 444.

31. Lindstad, L.J., Lo, G., Leivers, S., Lu, Z., Michalak, L., Pereira, G.V., Rohr, A.K., Martens, E.C., McKee, L.S., Louis, P., Duncan, S.H., Westereng, B., Pope, P.B., La Rosa, S.L. 2021. Human Gut Faecalibacterium prausnitzii Deploys a Highly Efficient Conserved System To Cross-Feed on beta-Mannan-Derived Oligosaccharides. mBio, 12(3), e0362820.

32. Liu, Y., Ma, J., Shi, R., Li, T., Yan, Q., Jiang, Z., Yang, S. 2021. Biochemical characterization of a β-N-acetylhexosaminidase from Catenibacterium mitsuokai suitable for the synthesis of lacto-N-triose II. Process Biochemistry, 102, 360–368.

33. Lombard, V., Golaconda Ramulu, H., Drula, E., Coutinho, P.M., Henrissat, B. 2014. The carbohydrate-active enzymes database (CAZy) in 2013. Nucleic Acids Research, 42(D1), D490–5.

34. Nath, A., Bhattacharjee, C., Chowdhury, R. 2013. Synthesis and separation of galacto-oligosaccharides using membrane bioreactor. Desalination, 316, 31–41.

35. NCBI. 2025a. PubChem Compound Summary for CID 6134, beta-Lactose. 2025 ed, National Center for Biotechnology Information. PubChem.

36. NCBI. 2025b. PubChem Compound Summary for CID 439242, Raffinose. 2025 ed, National Center for Biotechnology Information. PubChem.

37. Nekvasilova, P., Hovorkova, M., Meszaros, Z., Petraskova, L., Pelantova, H., Kren, V., Slamova, K., Bojarova, P. 2022. Engineered Glycosidases for the Synthesis of Analogs of Human Milk Oligosaccharides. International Journal of Molecular Sciences, 23(8).

38. Njoumi, S., Josephe Amiot, M., Rochette, I., Bellagha, S., Mouquet-Rivier, C. 2019. Soaking and cooking modify the alpha-galacto-oligosaccharide and dietary fibre content in five Mediterranean legumes. International Journal of Food Science and Nutrition, 70(5), 551–561.

39. Onyango, S.O., Beerens, K., Li, Q., Van Camp, J., Desmet, T., Van de Wiele, T. 2023. Glycosidic linkage of rare and new-to-nature disaccharides reshapes gut microbiota in vitro. Food Chemistry, 411, 135440.

40. Perna, V.N., Dehlholm, C., Meyer, A.S. 2021. Enzymatic production of 3’-sialyllactose in milk. Enzyme and Microbial Technology, 148, 109829.

41. Reddy, S.K., Bagenholm, V., Pudlo, N.A., Bouraoui, H., Koropatkin, N.M., Martens, E.C., Stalbrand, H. 2016. A beta-mannan utilization locus in Bacteroides ovatus involves a GH36 alpha-galactosidase active on galactomannans. FEBS Letters, 590(14), 2106–18.

42. Saldanha do Carmo, C., Knutsen, S.H., Malizia, G., Dessev, T., Geny, A., Zobel, H., Myhrer, K.S., Varela, P., Sahlstrøm, S. 2021. Meat analogues from a faba bean concentrate can be generated by high moisture extrusion. Future Foods, 3.

43. Saldanha do Carmo, C., Silventoinen-Veijalainen, P., Zobel, H., Holopainen-Mantila, U., Sahlstrøm, S., Knutsen, S.H. 2022. The effect of dehulling of yellow peas and faba beans on the distribution of carbohydrates upon dry fractionation. LWT, 163.

44. Sanz, M.L., Gibson, G.R., Rastall, R.A. 2005. Influence of disaccharide structure on prebiotic selectivity in vitro. Journal of Agricultural and Food Chemistry, 53(13), 5192–9.

45. Schober, I., Koblitz, J., Sarda Carbasse, J., Ebeling, C., Schmidt, M.L., Podstawka, A., Gupta, R., Ilangovan, V., Chamanara, J., Overmann, J., Reimer, L.C. 2025. BacDive in 2025: the core database for prokaryotic strain data. Nucleic Acids Research, 53(D1), D748–D756.

46. Segers, M.E., Lebeer, S. 2014. Towards a better understanding of Lactobacillus rhamnosus GG--host interactions. Microbial Cell Factories, 13 Suppl 1(Suppl 1), S7.

47. Shi, R., Ma, J., Yan, Q., Yang, S., Fan, Z., Jiang, Z. 2020. Biochemical characterization of a novel alpha-L-fucosidase from Pedobacter sp. and its application in synthesis of 3’-fucosyllactose and 2’-fucosyllactose. Applied Microbiology and Biotechnology, 104(13), 5813–5826.

48. Shin, Y.J., Woo, S.H., Jeong, H.M., Kim, J.S., Ko, D.S., Jeong, D.W., Lee, J.H., Shim, J.H. 2020. Characterization of novel alpha-galactosidase in glycohydrolase family 97 from Bacteroides thetaiotaomicron and its immobilization for industrial application. International Journal of Biological Macromolecules, 152, 727–734.

49. Swagerty, D.L., Jr., Walling, A.D., Klein, R.M. 2002. Lactose intolerance. American Family Physician, 65(9), 1845–50.

50. van Leeuwen, S.S., Kuipers, B.J.H., Dijkhuizen, L., Kamerling, J.P. 2014. (1)H NMR analysis of the lactose/beta-galactosidase-derived galacto-oligosaccharide components of Vivinal(R) GOS up to DP5. Carbohydrate Research, 400, 59–73.

51. van Zanten, G.C., Knudsen, A., Roytio, H., Forssten, S., Lawther, M., Blennow, A., Lahtinen, S.J., Jakobsen, M., Svensson, B., Jespersen, L. 2012. The effect of selected synbiotics on microbial composition and short-chain fatty acid production in a model system of the human colon. PLoS One, 7(10), e47212.

52. Varki, A., Cummings, R.D., Esko, J.D., Stanley, P., Hart, G.W., Aebi, M., Mohnen, D., Kinoshita, T., Packer, N.H., Prestegard, J.H., Schnaar, R.L., Seeberger, P.H. 2022. Essentials of Glycobiology. Cold Spring Harbor Laboratory Press.

53. Vogelsang-O’Dwyer, M., Petersen, I.L., Joehnke, M.S., Sorensen, J.C., Bez, J., Detzel, A., Busch, M., Krueger, M., O’Mahony, J.A., Arendt, E.K., Zannini, E. 2020. Comparison of Faba Bean Protein Ingredients Produced Using Dry Fractionation and Isoelectric Precipitation: Techno-Functional, Nutritional and Environmental Performance. Foods, 9(3).

54. Wang, K., Duan, F., Sun, T., Zhang, Y., Lu, L. 2024. Galactooligosaccharides: Synthesis, metabolism, bioactivities and food applications. Critical Reviews in Food Science and Nutrition, 64(18), 6160–6176.

55. Warmerdam, A., Benjamins, E., de Leeuw, T.F., Broekhuis, T.A., Boom, R.M., Janssen, A.E.M. 2014. Galacto-oligosaccharide production with immobilized β-galactosidase in a packed-bed reactor vs. free β-galactosidase in a batch reactor. Food and Bioproducts Processing, 92(4), 383–392.

56. Warmerdam, A., Paudel, E., Jia, W., Boom, R.M., Janssen, A.E. 2013. Characterization of beta-galactosidase isoforms from Bacillus circulans and their contribution to GOS production. Applied Biochemistry and Biotechnology, 170(2), 340–58.

57. Westereng, B., Kracun, S.K., Leivers, S., Arntzen, M.O., Aachmann, F.L., Eijsink, V.G.H. 2020. Synthesis of glycoconjugates utilizing the regioselectivity of a lytic polysaccharide monooxygenase. Scientific Reports, 10(1), 13197.

58. Wiull, K., Hagen, L.H., Roncevic, J., Westereng, B., Boysen, P., Eijsink, V.G.H., Mathiesen, G. 2024. Antigen surface display in two novel whole genome sequenced food grade strains, *Lactiplantibacillus pentosus* KW1 and KW2. Microbial Cell Factories, 23(1), 19.

59. Wong, S.Y., Hartel, R.W. 2014. Crystallization in lactose refining-a review. Journal of Food Science, 79(3), R257–72.

60. Yin, H., Pijning, T., Meng, X., Dijkhuizen, L., van Leeuwen, S.S. 2017a. Biochemical Characterization of the Functional Roles of Residues in the Active Site of the beta-Galactosidase from Bacillus circulans ATCC 31382. Biochemistry, 56(24), 3109–3118.

61. Yin, H., Pijning, T., Meng, X., Dijkhuizen, L., van Leeuwen, S.S. 2017b. Engineering of the Bacillus circulans beta-Galactosidase Product Specificity. Biochemistry, 56(5), 704–711.

62. Zeuner, B., Nyffenegger, C., Mikkelsen, J.D., Meyer, A.S. 2016. Thermostable beta-galactosidases for the synthesis of human milk oligosaccharides. New Biotechnology, 33(3), 355–60.

63. Zeuner, B., Teze, D., Muschiol, J., Meyer, A.S. 2019. Synthesis of Human Milk Oligosaccharides: Protein Engineering Strategies for Improved Enzymatic Transglycosylation. Molecules, 24(11).

64. Zhang, J., Song, G., Mei, Y., Li, R., Zhang, H., Liu, Y. 2019. Present status on removal of raffinose family oligosaccharides - a Review. Czech Journal of Food Sciences, 37(3), 141–154.

